# Visual, delay and oculomotor timing and tuning in macaque dorsal pulvinar during instructed and free choice memory saccades

**DOI:** 10.1101/2021.12.21.473504

**Authors:** Lukas Schneider, Adan-Ulises Dominguez-Vargas, Lydia Gibson, Melanie Wilke, Igor Kagan

**Affiliations:** Decision and Awareness Group, Cognitive Neuroscience Laboratory, German Primate Center, Leibniz Institute for Primate Research, Kellnerweg 4, Goettingen, 37077, Germany; Department of Cognitive Neurology, University Medical Center Göttingen, Robert-Koch-Str. 40, Goettingen, 37075, Germany; Département de Neurosciences, Faculté de Médecine, Université de Montréal, Québec, Canada; DFG Center for Nanoscale Microscopy & Molecular Physiology of the Brain (CNMPB), Robert-Koch-Str. 40, Göttingen, 37075, Germany; Leibniz ScienceCampus Primate Cognition, Kellnerweg 4, Goettingen, 37077, Germany

**Keywords:** eye movements, memory saccades, oculomotor decision, spatial choice selectivity

## Abstract

Causal perturbations suggest that the primate dorsal pulvinar (dPul) plays a crucial role in target selection and saccade planning, but its basic visuomotor neuronal properties are unclear. While some functional aspects of dPul and interconnected frontoparietal areas – e.g. ipsilesional choice bias after inactivation – are similar, it is unknown if dPul shares oculomotor properties of the cortical circuitry, in particular the delay and choice-related activity. We investigated such properties in macaque dPul during instructed and free-choice memory saccades. Most recorded units showed visual (16%), visuomotor (29%) or motor-related (35%) responses. Visual responses were mainly contralateral; motor-related responses were predominantly post-saccadic (64%) and showed weak contralateral bias. Pre-saccadic enhancement was infrequent (9-15%) – instead, activity was often suppressed during saccade planning (30%) and execution (19%). Surprisingly, only few units exhibited classical visuomotor patterns combining cue and continuous delay activity until the saccade or pre-saccadic ramping, and most spatially-selective neurons did not encode the upcoming decision during free-choice delay. Thus, in absence of a visible goal, the dorsal pulvinar has a limited role in prospective saccade planning, with patterns partially complementing its frontoparietal partners. Conversely, prevalent cue and post-saccadic responses imply the participation in integrating spatial goals with processing across saccades.

## Introduction

Visual information is crucial for guiding primate behavior. To gather this information, primates make saccadic eye movements towards locations of interest, allowing the fovea to obtain a better spatial resolution of these parts of the environment (Boi et al. 2017). Since eye movements are typically guided by visual inputs, it is not surprising that numerous cortical and subcortical regions show both visually-driven responses as well as saccade-related activity. To study visuomotor transformations underlying conversion of visual inputs into oculomotor actions, a classical memory-guided saccade task that allows separating visual, intervening delay and motor-related activity has been used extensively (Hikosaka and Wurtz 1983). With this task it has been shown that besides transient visually-evoked and motor-related responses, many neurons in oculomotor structures such as lateral intraparietal area (LIP), frontal eye fields (FFE), dorsal lateral prefrontal cortex (dlPFC), superior colliculus (SC), mediodorsal thalamus (MD) and central oculomotor thalamus show persistent delay activity. This activity links the visual input to an eventual motor output, and is considered a signature of many cognitive signals such as working memory, sustained attention, motor preparation, and evolving decision (Bruce and Goldberg 1985; Barash et al. 1991a; Funahashi et al. 1991; Basso and Wurtz 1998; Watanabe and Funahashi 2004a, 2007; Wyder et al. 2004, 2004; Lawrence et al. 2005; Kagan et al. 2010; Constantinidis et al. 2018).

Another important aspect of sensorimotor transformation concerns the continuity of visual experience and action guidance during a normal fixation-saccade cycle. Visual inputs enter the brain in eye-centered, retinotopic coordinates, but every saccade changes these representations, necessitating a mechanism for maintaining visual stability across saccades (Wurtz, Joiner, et al. 2011). It has been suggested that a corollary discharge (or efference copy) pathway involving SC, MD and FEF is one important contributor enabling anticipation of movement consequences and saccade suppression (Wurtz, Joiner, et al. 2011). The remapping of receptive fields around saccades in FEF and LIP is another related phenomenon thought to support visual stability (Bisley and Goldberg 2010; Mirpour and Bisley 2015). In addition to these prospective mechanisms manifesting prior to or during saccades, post-saccadic responses might also contribute to saccadic suppression, as well as to saccade updating and error processing (Zhou, Liu, et al. 2016).

Due to its widespread bidirectional connectivity to a host of visual and oculomotor cortical areas, the thalamic pulvinar is another likely candidate for mediating distributed mechanisms linking visual inputs to processing that spans across eye movements (Grieve et al. 2000; Guillery and Sherman 2002; Wilke, Turchi, et al. 2010; Berman and Wurtz 2011; Saalmann and Kastner 2015). Distinct pulvinar subdivisions might however contribute differently to these functions. Several studies have motivated a functional distinction between the ventral and the dorsal pulvinar aspects, based either on distinctive patterns of anatomical (Stepniewska et al. 1994; Kaas and Lyon 2007) and functional/effective connectivity (Arcaro et al. 2015, 2018; Kagan et al. 2021), or based on dissimilar functional properties within the same anatomically-defined subdivision, e.g. leading to further division of the lateral pulvinar into dorsal and ventral parts (Petersen et al. 1985; Baldwin et al. 2017). The ventral pulvinar (vPul, encompassing inferior pulvinar and ventral part of lateral pulvinar) is retinotopically organized and has strong connections to visual cortices (Kaas and Baldwin 2020). Ventral pulvinar neurons respond to contralateral visual stimuli but show weak or no pre-saccadic activity – most eye movement-related responses are peri- or post-saccadic (Petersen et al. 1985; Robinson et al. 1986; Berman and Wurtz 2011). In particular, the inferior pulvinar has been identified as a part of pathway from superficial layers of SC to cortical area MT that carries saccadic suppression signals (Berman and Wurtz 2011), while the SC - ventral lateral pulvinar pathway is involved in mediating the residual vision (blindsight) after V1 lesions (Kinoshita et al. 2019).

The dorsal pulvinar (dPul, encompassing medial pulvinar and dorsal part of lateral pulvinar) is reciprocally interconnected with posterior parietal and prefrontal cortices subserving attentional and sensorimotor functions, in particular LIP, area 7, FEF, dlPFC and posterior cingulate cortex (PCC) (Bos and Benevento 1975; Trojanowski and Jacobson 1976; Hardy and Lynch 1992; Gutierrez et al. 2000; Dean et al. 2004; Saalmann and Kastner 2011; Bridge et al. 2015; Bourgeois et al. 2020; Homman-Ludiye et al. 2020). Unlike the ventral pulvinar, the dorsal pulvinar does not follow clear retinotopic organization (Benevento and Miller 1981; Petersen et al. 1987; Benevento and Port 1995). Only few studies investigated visuomotor neuronal properties in the macaque dorsal pulvinar (in particular the medial part) using oculomotor tasks. Robinson, Petersen and colleagues (Petersen et al. 1985; Robinson et al. 1986) assessed responses to visually-guided saccades in the lateral part of the dorsal pulvinar (PLdm, denoted Pdm in the original studies). They showed that some Pdm neurons fired in association with eye movements, are crudely tuned for saccade direction and amplitude, and exhibit modulation of visual responses when stimuli are attended or serve as a saccade target. Another study showed that the visual responsivity to stimuli in the receptive field is reduced in the lateral dorsal pulvinar when monkeys attend to the central fixation, further suggesting the importance of behavioral relevance and involvement of Pdm/PLdm in selective visual attention (Bender and Youakim 2001).

To our knowledge, only one study prior to our work investigated neurons in the dorsal pulvinar with the memory-guided saccade task (Benevento and Port 1995). The main focus of this study was on color and pattern processing; the comparison of visually-guided and memory-guided task responses in a small sample indicated that stimulus-unrelated activity was post-saccadic. Surprisingly, in this study visual target-related responses were reported during visually-guided but not memory-guided saccades, and the delay activity was not addressed. The authors suggested that only when the stimulus serves as an *immediate* saccade target the visual response takes place, although it is not the pattern typically observed in the frontoparietal cortical areas. Finally, we recently have shown that dPul neurons carry information about static eye position and combine spatial encoding in eye-centered and nonretinocentric coordinates, potentially contributing to coordinate frame transformations (Schneider et al. 2019).

In contrast to paucity of single neuron studies on oculomotor processing and target selection in the dorsal pulvinar, more is known about consequences of its causal perturbation. While primary sensory and oculomotor functions are largely spared (Ungerleider and Christensen 1977; Bender and Butter 1987; Bender and Baizer 1990; Wilke et al. 2013), dPul inactivation or microstimulation affects saccadic reaction times and biases hemifield-specific spatial exploration and target selection, especially in conditions of a free choice (Wilke, Turchi, et al. 2010; Wilke et al. 2013; Dominguez-Vargas et al. 2017). The inactivation of the lateral part of the dorsal pulvinar also impairs contralateral attentional orienting (Petersen et al. 1987; Desimone et al. 1990) and perceptual decision confidence (Komura et al. 2013). It is not clear however whether the transformation from visual processing to motor actions is implemented already within the dPul at the level of individual neurons, or these transformations are only taking place in the interconnected cortical circuitry. Furthermore, it is not known if dPul neurons show delay period or pre-saccadic spatial choice selectivity, similar to frontoparietal cortex (Coe et al. 2002; Watanabe and Funahashi 2007) and other higher-order thalamic nuclei such as central and mediodorsal thalamus (Watanabe and Funahashi 2004a; Wyder et al. 2004; Costello et al. 2015).

In a recent electrical microstimulation study of choice behavior in the dorsal and ventral pulvinar (Dominguez-Vargas et al. 2017), we provided initial electrophysiological assessment of microstimulation sites in dPul (mostly in the medial pulvinar). A detailed characterization of dPul with a classical memory-guided saccade task, an invaluable tool for understanding visuomotor processing, is however lacking. To fill this gap, here we assessed the temporal evolution of enhancement, suppression and spatial tuning of visual and motor related activity, and quantified the relationship between visual and motor responses. Furthermore, we addressed the question how pulvinar activity might relate to prospective visuomotor transformations during motor preparation by systematically analyzing the delay period activity. Given the strong effect of pulvinar perturbation on spatial choices, we also investigated whether dPul neurons signal upcoming decisions in free-choice two-target trials.

## Materials and Methods

All experimental procedures were conducted in accordance with the European Directive 2010/63/EU, the corresponding German law governing animal welfare, and German Primate Center institutional guidelines. The procedures were approved by the responsible government agency (“Niedersächsisches Landesamt für Verbraucherschutz und Lebensmittelsicherheit” - Lower Saxony State Office for Consumer Protection and Food Safety, LAVES, Oldenburg, Germany; permit 3392-42502-04-13/1100). The animals were pair- or group-housed in facilities of the German Primate Center in accordance with all applicable German and European regulations. The facility provides the animals with an enriched environment, including a multitude of toys and wooden structures (Calapai et al. 2017; Berger et al. 2018), natural as well as artificial light and access to outdoor space, exceeding the size requirements of European regulations, and a rich diet including primate biscuits, fruit and vegetables.

### Animal preparation

Two adult male rhesus monkeys (*Macaca mulatta*) C and L weighing 8 and 9 kg respectively were used. Monkeys were pair-housed with other males in two enclosures (23.3 m^3^ and 21.3 m^3^) conjoined by an overhead tunnel. In an initial surgery, monkeys were implanted with a magnetic resonance imaging (MRI) compatible polyetheretherketone (PEEK) headpost embedded in a bone cement headcap (Palacos with Gentamicin, BioMet, USA) anchored by ceramic screws (Rogue Research, Canada), under general anesthesia and aseptic conditions. MR-visible markers were embedded in the headcap to aid the planning of the chamber in stereotaxic space with the MR-guided stereotaxic navigation software Planner (https://github.com/shayo/Planner) (Ohayon and Tsao 2012). A separate surgery was performed to implant PEEK MRI-compatible chamber(s) (inside diameter 22 mm) allowing access to the pulvinar (Monkey C, right hemisphere: center at 0.5A / 14.5R mm, tilted −11P / 27R degrees; Monkey L, right hemisphere: center at −3.12P / 20.2R mm, tilted −18P/37R degrees; Monkey L, left hemisphere: center at −3P/20L, tilted −18P/−38L; the coordinates are relative to stereotaxic zero, A – anterior, P – posterior, L – left, R – right; the tilts are relative to the vertical axis normal to stereotaxic plane: P – posterior, top of the chamber tilted towards the back of the head, L/R – left/right, top of the chamber tilted towards the corresponding ear). After confirming chamber positioning with a post-surgical MRI, a partial craniotomy was made inside the chamber. The exposed dura was covered with a silicone elastomer (Kwik-sil, World Precision Instruments, USA) to reduce the granulation tissue growth and dura thickening.

### MRI imaging

Monkeys were scanned in a 3T MRI scanner (Siemens Magnetom TIM Trio). Full-head T1-weighted scans (3D magnetization-prepared rapid gradient-echo, MPRAGE, 0.5 mm isometric) were acquired before and after chamber implantation, in awake (monkey C) or anaesthetized (monkey L) state, using either built-in gradient body transmit coil and custom single loop receive coil, or custom single loop transmit and 4-channel receive coil (Windmiller Kolster Scientific, USA).

In addition to pre- and post-implantation scans, similar T1-weighted scans as well as T2-weighted (rapid acquisition with relaxation enhancement, RARE, 0.25 mm in plane resolution, 1 mm slice thickness) scans were periodically acquired during the course of experiments, either in awake (monkey C) or sedated (monkey L) state, to confirm electrode positioning. These images were acquired with the chamber and a custom-made MR-compatible polyetherimide (Ultem) grid (0.8 mm hole spacing, 0.45 mm hole diameter) filled with gadolinium (Magnevist, Bayer, Germany)/saline solution (proportion 1:200), with tungsten rods inserted in predefined grid locations, for alignment purposes. The electrode penetration trajectory and the recording depth was visualized using platinum-iridium electrodes (FHC Inc., USA, 100 mm length platinum-iridium 125 μm thick core, initial 2 cm glass-coated, total thickness of 230 μm including polyamide tubing coating, UEPLEFSS (UEIK1)), advanced by a MRI-compatible custom-made plastic XYZ manipulator drive through the corresponding grid hole. For the dura penetration, the electrode was protected by a custom-made MRI-compatible fused silica guide tube (320 µm inner diameter, 430 µm outer diameter, Polymicro Technologies, USA). A stopper (530 µm inner diameter, 665 µm outer diameter, 23 gauge MicroFil, World Precision Instruments, USA) ensured that the guide tube only penetrated the dura and minimally the cortex below. Prior to penetration, the electrode tip was aligned to the guide tube tip and was held in place by a drop of melted Vaseline. The guide tube was filled with sterile silicone oil prior to electrode insertion, to ensure smooth electrode travel and to prevent the backflow of cerebrospinal fluid. T1- and T2-weighted scans were co-registered and transformed into “chamber normal” (aligned to the chamber vertical axis) and to anterior commissure – posterior commissure (AC-PC) space for electrode targeting and visualization in BrainVoyagerQX (Brain Innovation, Netherlands) and Planner (Ohayon and Tsao 2012; Kagan et al. 2021).

### Electrophysiological recordings

Dorsal pulvinar neuronal activity was recorded with up to four individually-movable single platinum-tungsten (95%-5%) quartz glass-insulated electrodes with impedance ranging from 1 MΩ to 1.9 MΩ for monkey C and 1 MΩ to 3.5 MΩ for monkey L, using a chamber-mounted 5-channel Mini Matrix microdrive (Thomas Recording, Germany). Single custom-made stainless steel guide tubes (27 gauge), made from Spinocan spinal needle (B. Braun Melsungen AG, Germany) filled with silicone oil (Thomas Recording), with a metal funnel attached to the microdrive nozzle were used to protect electrodes during grid insertion and dura penetration. A reference tungsten rod or a silver wire was placed in the chamber filled with saline, and was connected to the chassis of the drive. Neuronal signals were amplified (×20 headstage, Thomas Recording; ×5, 128 or 32 channel PZ2 preamplifier, Tucker-Davis Technologies, USA), digitized at 24 kHz and 16-bit resolution, and sent via fiber optics to an RZ2 BioAmp Processor (Tucker-Davis Technologies, USA) for online filtering (300 – 5000 Hz bandpass), display and storage on a hard drive together with behavioral and timing data streams.

### Behavioral tasks

Monkeys sat in a dark room in custom-made primate chairs with their heads restrained 30 cm away from a 27’’ LED display (60 Hz refresh rate, model HN274H, Acer Inc. USA), covering a range of 84×53 visual degrees for monkey C and 91×59 for monkey L. The gaze position of the right eye was monitored at 220 Hz using an MCU02 ViewPoint infrared eyetracker (Arrington Research Inc. USA). Eye position was calibrated using linear gains and offsets for horizontal and vertical axes, using visually-guided saccades, and recalibrated using MATLAB (The MathWorks, Inc. USA) geometric transformation function fitgeotrans.m with 2^nd^ degree polynomial fit. All stimulus presentation and behavioral control tasks were programmed in MATLAB and the Psychophysics Toolbox (Brainard 1997), using a custom monkeypsych toolbox (https://github.com/dagdpz/monkeypsych).

#### Memory-guided saccade task

A trial started with the onset of the fixation point of 1° diameter. After the gaze fixation was acquired and held within a 5° radius for 500 ms, either one peripheral 1° diameter cue (instructed trials) or two peripheral cues (choice trials) were displayed for 300 ms at the location(s) signaling the upcoming saccade target(s). Cues were presented in the left and/or right side(s) of the fixation point, at 12° or 24° absolute eccentricity, with six potential directions relative to the horizontal axis: circular angles counterclockwise 0°, 20°, 160°, 180°, 200°, 340° (resulting in 0°, 4.1°, −4.1°, 8.2° and −8.2° vertical eccentricity). Monkeys were required to maintain fixation throughout the cue period and the subsequent memory period (1000 ms), after which the central fixation point disappeared, prompting monkeys to saccade to the remembered instructed location, or to choose one of the two cued locations, within 500 ms. The fixation offset will be referred to as the Go signal. After a saccade to and fixation inside a 5° radius window around the remembered target location for 100 or 200 ms the selected target became visible, and after additional 500 ms of peripheral fixation the trial was completed and a liquid reward was dispensed after a delay of 200 ms. Instructed trials (50%) were interleaved with choice trials (50%), and trial types and target locations were pseudo-randomized. In choice trials the two cues were always presented simultaneously at the same height and provided equal reward. The inter-trial interval for both successful and unsuccessful trials was 2000 or 2500 ms (**Figure 1A**).

**Figure 1.**
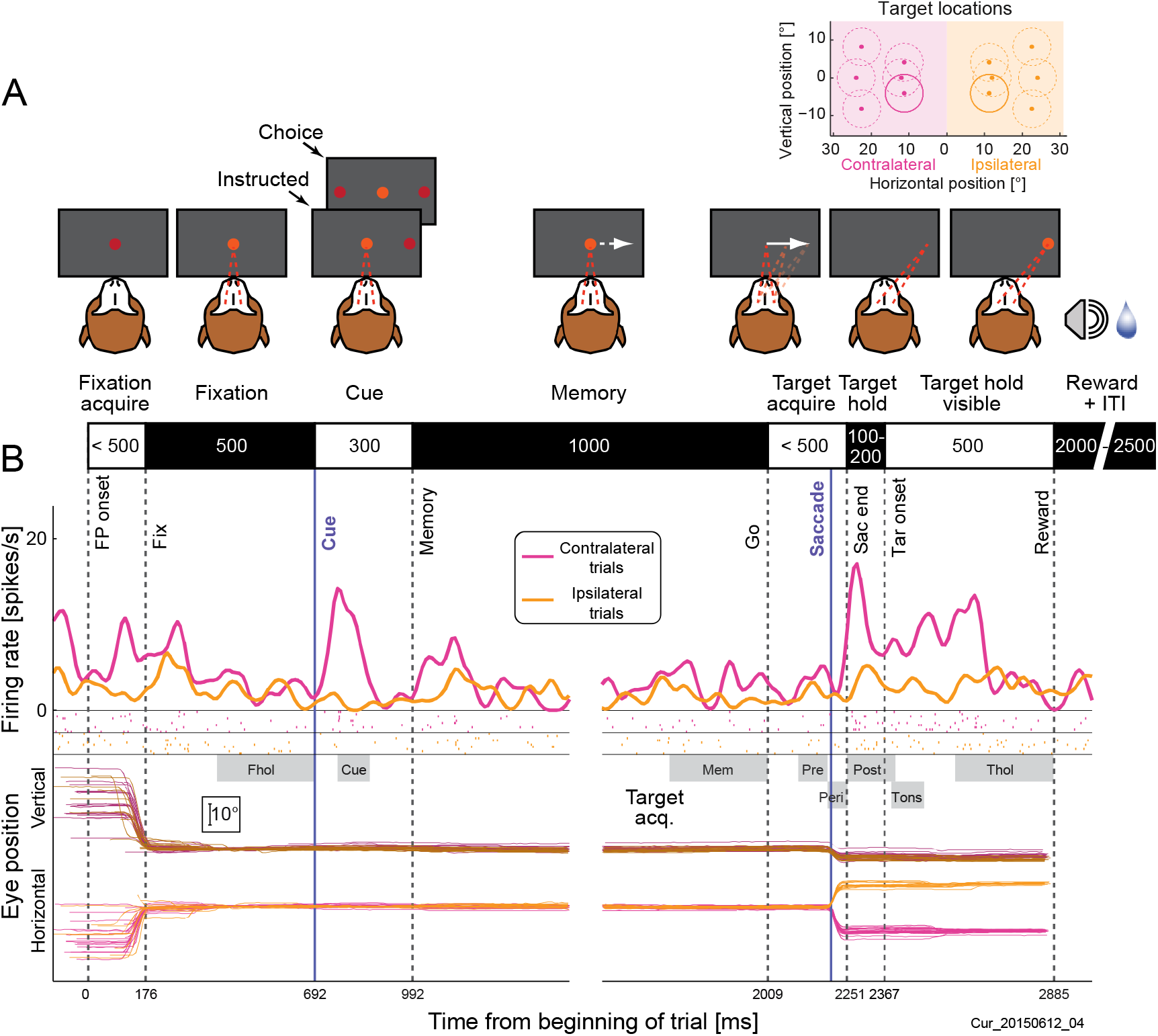
Memory-guided saccade task and example neuron activity. **A:** Task layout. Monkeys had to fixate in the center while one or two cues were flashed peripherally. After a 1 s memory period monkeys had to make a saccade to (one of the) previously cued location(s), and fixate this location until and after the target became visible in order to receive reward. In instructed trials, there was one target; in choice trials - two targets. The two targets in choice trials were always presented at the same height and eccentricity and provided equal reward. Inset on top right shows all target positions and fixation windows; solid fixation windows indicate two targets in trials shown in B. **B:** Raster plots and spike density functions for two example positions, a contralateral (magenta) and the opposite ipsilateral (orange) position. Corresponding eye traces are below (darker for vertical eye position, brighter for horizontal eye position). Vertical lines indicate average onset of events across all trials for this recording block (including other target positions): fixation point onset (FP onset), monkey acquiring fixation (Fix), the cue onset, the cue offset and beginning of the memory period, the offset of the central fixation point (Go), the saccade onset, monkey acquiring the invisible target location (Sac end), the onset of the visible peripheral target (Tar onset), and the end of the trial (Reward). Discontinuous traces indicate two different alignments to cue onset and onset events (purple lines). Grey boxes mark analyzed epochs: fixation hold (Fhol), cue onset (Cue), memory (Mem), pre-saccadic (Pre), peri-saccadic (Peri), post-saccadic (Post), Target onset (Tons), and target hold (Thol) (see **Materials and Methods**).

#### Visually-guided saccade task

In addition to the memory-guided saccade task, monkeys also performed visually-guided saccades in separate blocks in most of the sessions. In this task, after the initial fixation period the fixation point disappeared and the visual target(s) appeared at the same time, serving as the Go signal. Monkeys had to saccade towards the visible peripheral target and keep fixating it for 500 ms to obtain a liquid reward.

Target sizes, colors and locations were the same as for memory-guided saccades, and trial types and target locations were pseudo-randomized.

### Data analysis

#### Saccade definition

Instantaneous saccade velocity was calculated sample by sample as the square root of the sum of squared interpolated (220 Hz to 1 kHz) and smoothed (12 ms moving average rectangular window) horizontal and vertical eye position traces, and then smoothed again (12 ms moving average rectangular window). Saccade onset was defined as the first eye position change after the go signal that exceeded a velocity threshold of 200°/s; saccade end was defined as the first eye position change below 50°/s after saccade onset. In instructed memory-guided trials, average saccade latency across sessions was 199 ± 2 ms (212 ± 1 ms for monkey C, 191 ± 1 ms for monkey L); average saccade duration 45.6 ± 0.2 ms (46.8 ± 0.2 and 44.9 ± 0.2), average maximum saccade velocity was 622 ± 4°/s (594 ± 3°/s and 640 ± 4°/s). For corrective saccades (small re-fixation saccades after target reappearance), 20°/s and 10°/s were used as onset and offset velocity thresholds. Corrective saccades were common in both monkeys (90 ± 2% and 92 ± 1% of all trials), with an average latency of 125 ± 3 ms (138 ± 5 ms and 116 ± 2 ms) after target reappearance and average amplitude of 1.08 ± 0.02° (1.00 ± 0.03° and 1.13 ± 0.02°). All values in this section are session means ± SE, values in parentheses for monkey C and for monkey L separately. For details see **Supplementary Table S1**.

#### Datasets and unit selection criteria

All voltage drops that crossed an online visually determined threshold were defined as potential spikes and recorded on the hard drive. Spike sorting was done in Offline Sorter v.3.3.5 (Plexon, USA), using a waveform template algorithm, after manually defining templates by clustering in principle component space using first three principle components, as well as the recording time axis. Based on inspection of spike shapes, principle component cluster shapes and inter-spike intervals, clusters were categorized as single units, multi-units, and rejected “noise”.

In total, 416 single and multi-units were recorded in the dorsal pulvinar (dPul) in 50 sessions during the memory-guided saccade task (monkey C; right hemisphere: 235, monkey L; right hemisphere: 123; left hemisphere: 58). Out of these, 371 units (200 - monkey C right dPul, 115 - monkey L right dPul, 56 - monkey L left dPul) fulfilled analysis selection criteria (stable discriminability across time and acceptable signal-to-noise ratio – assessed by inspection). Out of these 371 units, 322 units (174 monkey C right dPul, 98 monkey L right dPul, 50 monkey L left dPul) were recorded for at least 4 successful instructed trials for each of the 12 targets and were used for further analysis. The median (across units) minimal number of trials per instructed target position was 9 trials (with a range from 4 to 29 trials). Most units (251 out of 322) were recorded for at least 9 trials per target position. The median (across units) average number of trials per position was 18 with a range from 6 to 39 trials.

Out of 322 unit accepted for the analysis, 249 units (126 monkey C right dPul, 82 monkey L right dPul, 41 monkey L left dPul) were categorized as well-isolated stable single units. Out of these 249 units, 245 units were also recorded in at least 4 choice trials for each hemifield and used for spatial choice selectivity analysis that only considered single units. All other analyses were performed on single and multi-units together.

The mean firing rate of 249 well-isolated stable single units was 13.4 spikes/s (median: 9.0, SD: 11.7), as compared to 17.5 spikes/s (median: 11.2, SD: 14.9) for the 73 remaining multi-units, and 14.5 spikes/s across all 322 units (median: 9.7, SD: 12.8).

Out of 322 units analyzed for the memory-guided task, 195 units (101 monkey C right dPul, 54 monkey L right dPul, 40 monkey L left dPul) were also recorded during the visually-guided saccade task. Out of these 195 units, 174 (88 monkey C right dPul, 46 monkey L right dPul, 40 monkey L left dPul) were recorded for at least 4 successful instructed trials for each of the 12 targets. These 174 units were used for comparison between the two tasks.

#### Epoch definitions and modulation

For each trial, and each epoch of interest, firing rates were computed by counting spikes within the epoch and dividing by the epoch duration. The following epochs were analyzed: inter-trial interval (400 ms to 100 ms before the onset of the central fixation point, corresponding to the fixation acquire), fixation hold (last 300 ms of central fixation), cue onset (70 ms to 170 ms after cue onset), memory (last 300 ms of the memory period), pre-saccadic (100 to 10 ms before saccade onset), peri-saccadic (10 ms before to 50 ms after saccade onset), post-saccadic (first 150 ms after acquiring the invisible peripheral target), target onset (20 ms to 120 ms after the target became visible), and target hold (last 300 ms of fixating the peripheral target) (**Figure 1B**).

The cue onset epoch was defined according to findings in the response timing analysis as 100 ms around the enhancement peak, see **Figure 2D**. We did not base our post-saccadic epoch on such analysis, for two reasons. First, the onset of post-saccadic modulation was not as synchronized across units as the cue response (*cf*. **Figure 2F**). Second, in some sessions the target became visible already 100 ms after saccade offset, which could lead to confounding visual responses, giving only short interval for a purely post-saccadic epoch. Instead, we defined the post-saccadic epoch as the first 150 ms after saccade offset. In those cases, when the target appeared already 100 ms after the saccade offset, this epoch would not be contaminated by visual response to the target onset because visual latency of most pulvinar neurons was >50 ms.

**Figure 2.**
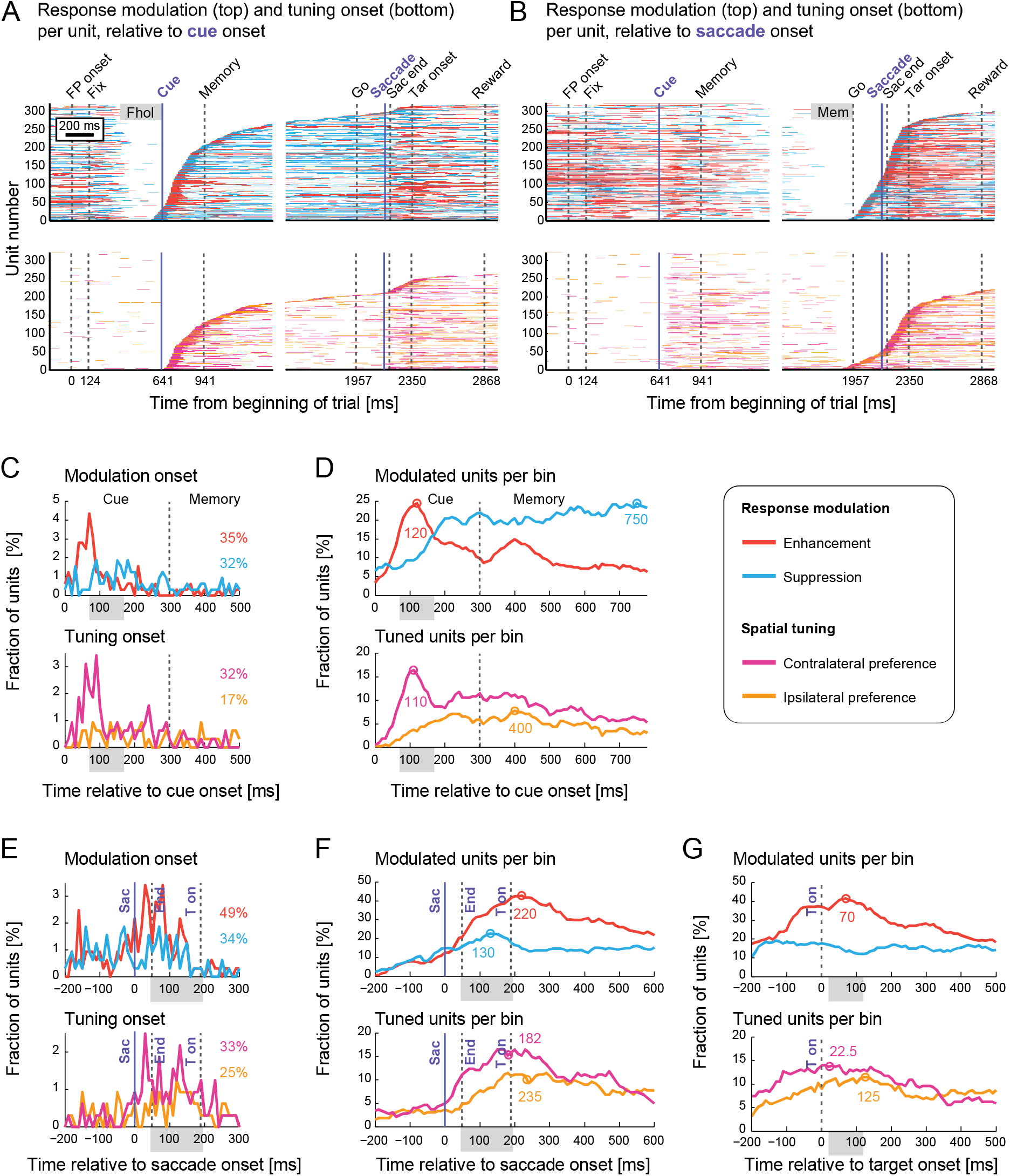
Response modulation and spatial tuning onsets. Bin-by-bin significance of modulation for all 332 units, in 10 ms bins. **A:** Non-spatial specific response modulation (Top) and hemifield preference (Bottom) across time for all units recorded, ordered by modulation onset time relative to cue onset. Each horizontal line represents one unit. Non-spatial specific modulation is shown as enhancement (red) or suppression (blue) relative to the baseline epoch fixation hold (Fhol), indicated by a gray box. Hemifield preference is shown as contralateral preference (magenta) or ipsilateral preference (orange). **B:** Same as A, ordered by modulation onset relative to saccade onset. Here, the memory epoch (Mem) was used as the baseline epoch for non-spatial specific modulation. **C:** Fraction of units showing modulation and tuning onset for each 10 ms bin relative to cue onset. Numbers indicate the sum across all bins in the displayed time range. The grey box denotes the cue onset epoch, the dotted line denotes the cue offset. **D:** Fraction of units showing modulation and tuning for each 10 ms bin, relative to cue onset. Colored circles and numbers show the time points corresponding to the maximum number of modulated or tuned units. In case there were several maxima with the same amount of units, the respective time points were averaged (as in F, lower panel). **E:** Same as C, relative to saccade onset. The grey box denotes the post-saccadic epoch. Vertical lines show the saccade onset (purple), the saccade end and the onset of the visible target (dotted lines). **F:** Same as D, relative to saccade onset. **G:** Same as D, relative to target onset. The grey box denotes the target onset epoch.

To compare firing rates in the target onset/pre-saccadic epoch ‘target-pre-sac’ of the visually-guided saccade task (40-140 ms after target onset/Go signal) to firing rates in the memory-guided saccade task, two additional corresponding epochs were defined for the memory-guided task: ‘cue onset 2’ (40-140 ms after cue onset) and ‘pre-saccadic 2’ (130-30 ms before saccade onset).

For most analyses, data from 6 left and 6 right hemifield targets were combined. For each unit, spatial tuning in each epoch was determined by unpaired t-tests comparing firing rates in ipsilateral trials to firing rates in contralateral trials. The hemifield with the higher firing rate was marked, if there was a significant difference. Enhancement or suppression of neuronal activity in each epoch was defined by t-tests comparing firing rates to a respective preceding baseline epoch, independently for ipsilateral and contralateral trials. For fixation hold epoch inter-trial interval served as baseline, for cue onset and memory epochs (as well as ‘cue onset 2’ and ‘pre-saccadic 2’ for comparison with visually-guided task), the fixation hold epoch served as baseline. The memory epoch served as baseline for all subsequent epochs. Enhancement or suppression was reported, if either ipsilateral, contralateral, or both types of trials showed significant difference to fixation baseline. In rare cases where one hemifield would show a significant enhancement, while the other hemifield showed suppression, the unit was reported to have bidirectional response.

Units that showed enhancement or suppression for at least one hemifield in both cue onset and post-saccadic epochs were classified as visuomotor units, whereas units that only showed enhancement or suppression in one of the two epochs were classified as visual (only enhancement or suppression in cue onset) and motor (only enhancement or suppression in post-saccadic). Visuomotor index was defined independently for ipsilateral and contralateral trials as (M-V)/(M+V), where M is the absolute average firing rate change in post-saccadic relative to the memory epoch, and V is the absolute average firing rate change in cue onset relative to fixation hold (Lawrence et al. 2005).

In addition to the visuomotor index described above, we also used visuomotor categories derived using enhancement or suppression in cue onset and pre-saccadic (as opposed to post-saccadic) epochs relative to fixation hold. This was done to compare the results to previously used visuomotor categorizations (Bruce and Goldberg 1985; Wyder et al. 2003).

To evaluate how many neurons exhibited task-related modulation, for each unit we applied a two-way ANOVA [hemifield × epoch] on firing rates across all successful trials, using hemifield of the target position and epoch as factors for determining a main effect of epoch, a main effect of the hemifield, and interaction between the two factors.

#### Response modulation onset and tuning onset analysis

To evaluate the time of response modulation onset for each unit and further define the cue onset epoch, spike density functions for each trial were derived by convolution of the discrete spike arrival times with a Gaussian kernel (standard deviation 20 ms). For three alignments (to cue onset, saccade onset and target onset), spike densities for each unit and each bin was compared to the average firing rate in the respective baseline epoch (fixation hold for cue alignment and memory for saccade and target alignment) using paired t-tests across all trials to evaluate bin-by-bin modulation. Significant increases were reported as enhancement and significant decreases as suppression. Similarly, for each unit and each bin, the spike densities of contralateral trials and ipsilateral trials were compared using unpaired t-tests. Significantly higher spike density for contralateral trials was reported as contralateral preference and significantly higher spike density for ipsilateral trials as ipsilateral preference. To avoid overestimation of significance due to multi-comparison, significance in less than five consecutive bins was discarded. For each unit and each type of modulation, the modulation onset was taken as the first bin (out of at least 5 consecutive bins) after certain time point that showed significance. For modulation onset relative to cue onset, the first significant bin after cue onset was taken and for modulation onset relative to the saccade the first significant bin after 200 ms before saccade onset was taken. Additionally, we counted the number of units which showed significant modulation in each bin and calculated the time in which most units were modulated. Based on the finding that most units showed enhancement 120 ms after cue onset (*cf*. **Figure 2D**), the cue onset epoch was selected as 50 ms before to 50 ms after that maximum.

#### Response fields

To estimate the center and the extent of visual and post-saccadic response fields (RFs), we fitted 2D Gaussian distributions on the response strength averages per target position, weighted by inverse of response variation per position, in cue onset (Cue) and post-saccadic (Post) epoch. Response strengths were calculated relative to the respective baseline, by subtracting firing rates in fixation hold (Fhol) for visual cue responses and firing rates in the memory epoch (Mem) for post-saccadic responses. This analysis was only performed on units that showed enhancement or suppression for at least one hemifield in the respective epoch. The fitting of RFs was performed in two steps: first, one Gaussian RF zone was fitted, allowing either a positive or a negative peak. The second Gaussian RF zone was fitted allowing only the opposite sign of the peak, and not allowing overlap with the first zone. Significant fits were reported if the absolute residuals of at least one of the RF zones were significantly smaller than the absolute residuals to the zero plane fit (p<0.05, one-tailed unpaired t-test). A response field with two zones was reported if absolute residuals were significantly smaller when using both RF zone fits than when only using the first RF zone fit (p<0.05, one-tailed unpaired t-test).

Six fitting parameters for each Gaussian response zone were determined using an iterative least squares method (400 iterations), allowing elliptic response zones with peaks (or troughs) at the center. The fitting parameters were (1) the response strength in the center of zone, (2) horizontal and (3) vertical location of the center of the response zone, (4 and 5) standard deviations describing semi-minor and semi-major axes, and (6) an angle of rotation of these axes. Importantly, the response zone centers were always kept within the dimensions of the target array (−24 to +24 degrees horizontally and (−8 to +8 degrees vertically). The response strength was bounded by −150% and 150% of the original maximum response strength, standard deviations were bounded by 1.5° (a quarter of the maximum horizontal target distance) and 12° (a quarter of the horizontal extend of the target array), and the rotation was unbounded.

The size of each elliptic zone was defined by two standard deviations in each direction (semi-minor and semi-major axes). A radius r approximating response zone size was calculated by taking the square root of the product of the two axes of the elliptic zones. This way, πr^2^ always matches with the area covered by the elliptic response zone. Response zone size is reported as diameter, 2r. For units with one response zone, response field size was defined equal to response zone size. For units with two response zones, we reported the size of the larger response zone.

#### Peri-stimulus time histograms (PSTHs)

To calculate population PSTHs, spike density functions of each trial were either baseline-corrected by subtracting the average ongoing firing rate in the inter-trial-interval epoch or normalized by dividing by the average firing rate (across all trials) in the fixation hold epoch. Average responses for each unit were then derived by averaging the normalized spike density for each unit across all trials for the respective condition. Means and SE of these baseline-corrected and averaged spike densities across units of a given sub-population were calculated to display population responses.

#### Choice selectivity

To see if information about the upcoming target selection could be derived from spiking activity, we correlated instructed and choice firing rate differences between contralateral and ipsilateral trials using Pearson’s correlation. For this analysis, units were considered only if at least four choice trials to each hemifield were recorded. The contralaterality index in instructed and choice trials was calculated as CI = (firing rate _contra_ – firing rate _ipsi_ / (firing rate _contra_ + firing rate _ipsi_). For comparing the absolute strength of spatial selectivity in instructed vs. choice trials, we subtracted the absolute CI_instr_ and CI_choice_ values and tested the significance of difference using paired Wilcoxon’s signed rank test.

## Results

### Response timing and tuning in memory-guided saccade task

We recorded dorsal pulvinar neuronal activity in two monkeys performing a memory-guided saccade task. Trials always started with initial fixation in the center of the screen followed by a cue or two cues, memory delay and saccade to one out of 12 peripheral instructed or chosen locations (**Figure 1A** and **Materials and Methods**).

The activity of one example unit and the eye traces for two target positions are shown in **Figure 1B**. This unit shows an enhanced visual response after the cue presentation at contralateral positions as well as a contralateral post-saccadic response which cannot be attributed to a target-related visual response, because it occurs before the onset of the visual target (Tar onset).

To evaluate how population activity of all 322 units selected for the analysis developed over trial time (and to validate the timing of analysis epochs), we estimated significant enhancement or suppression (combined across both hemifields) as well as hemifield preference for each unit in each 10 ms bin (**Figure 2**, see also **Materials and Methods**). In **Figure 2A** units are ordered by the onset of significant modulation relative to cue onset. The fixation hold epoch before cue onset was used as baseline for calculating enhancement and suppression (**Figure 2A**, top). This analysis showed a time-locked enhancement shortly after the cue onset and a predominance of suppression during the memory period (**Figure 2A**, top) combined with contralateral cue preference shortly after the cue onset and an overall lack of spatial preference during the memory epoch (**Figure 2A**, bottom). In **Figure 2B** units are ordered by the onset of significant modulation relative to saccade onset. Because of the frequent differences in firing rate between initial fixation and memory periods, we used the memory epoch as baseline for calculating enhancement and suppression around the saccade (**Figure 2B**, top). Around the saccade we observed predominantly post-saccadic enhancement (**Figure 2B**, top) and a bias to contralateral hemifield (**Figure 2B**, bottom).

A more detailed picture can be obtained by looking at the distribution of response modulation and spatial tuning *onsets* relative to cue (**Figure 2C**), and the number of units which showed respective property at each 10 ms bin after cue onset (**Figure 2D**). While onset of enhancement and contralateral preference typically started at around 50 ms after cue onset, the timing of suppression and ipsilateral preference was more dispersed (**Figure 2C,D** *cf*. blue vs. red and orange vs. magenta). Enhancement (independent of spatial preference) was most common at 120 ms after cue onset, close to the time bin where the maximum number of units showed contralateral preference (110 ms after cue onset). The same analysis for modulation relative to saccade onset (**Figure 2E-F**) showed that pre-saccadic and peri-saccadic modulation was rare compared to post-saccadic modulation, and that enhancement and contralateral preference were widely dispersed in the post-saccadic period, reaching maximum at 220 ms after saccade onset (or 170 ms after saccade end).

Many units showed enhancement and spatial preference well before the target onset (**Figure 2G**), further confirming that post-saccadic responses are mainly not due to direct visual stimulation. Additionally, out of 34 units (11%) with a significant hemifield preference both for cue and post-saccadic responses, only 11 units preferred the opposing hemifields in the two epochs, while 23 units showed the same hemifield preference in both epochs. This finding rules out the possibility that the post-saccadic response in those units is driven by increased luminance flux from the monitor background backlight when the peripheral RF is shifted by the saccade from the space beyond the edge of the monitor (covering 90° visual angle in the horizontal dimension) onto the monitor – because if this were the case, the cue and the post-saccadic responses should have the opposite preference.

Interestingly, the fraction of units that showed enhancement after the target onset reached the maximum at 70 ms (**Figure 2G**), earlier then for cue responses (120 ms). This might indicate a difference in timing of visual responses for peripheral (cue) and foveal (target) stimuli. The post-saccadic peak was however wide and likely results from confluence of post-saccadic and visual target onset-related signals.

To summarize, the predominant response patterns were contralateral cue enhancement (peaking at 120 ms after the cue onset), suppression throughout the entire memory period, post-saccadic enhancement (wide peak at 220 ms) and peri/post-saccadic suppression (wide peak at 130 ms), often accompanied by post-saccadic hemifield preference (more often contralateral than ipsilateral). **Figure 3** shows the summary of response modulation, spatial and hemifield tuning in all task-relevant epochs in instructed trials. For each unit and analysis epoch, we evaluated significant enhancement or suppression (now per hemifield, as compared to the preceding response modulation timecourse analysis in **Figure 2**, done across the two hemifields combined), and hemifield preference. If there was no significant hemifield preference, we also tested one-way ANOVA across 12 target positions, to account for units with upper/lower selectivity or idiosyncratic tuning profile. The majority of units (89%) showed task-related modulation in at least one of the analyzed epochs. Contralateral hemifield preference was more common than ipsilateral preference in the cue onset epoch (24% vs. 6%). Importantly, most units with contralateral cue preference (51/76) showed the enhancement. The number of units preferring contralateral or ipsilateral hemifield became more equalized in subsequent epochs (with a remaining weak contralateral bias), and hemifield preference was less frequent in memory (16%), pre-saccadic (11%) and peri-saccadic epoch (14%) as compared to post-saccadic (30%), target onset (34%) and target hold (33%) epochs. Enhancement was more common than suppression during the cue period (30% vs. 13%), while the opposite was true for the memory period (16% vs. 30%) as well as before and during the saccade (9% vs. 14% in pre-saccadic and 13% vs. 19% in peri-saccadic epochs). After the saccade, activity was typically enhanced (40% enhancement vs. 21% suppression in post-saccadic, 46% vs. 19% in target onset and 36% vs. 19% in target hold epochs). For details on how interrelated were enhancement/suppression and spatial tuning patterns, see **Supplementary Figure S1**.

**Figure 3.**
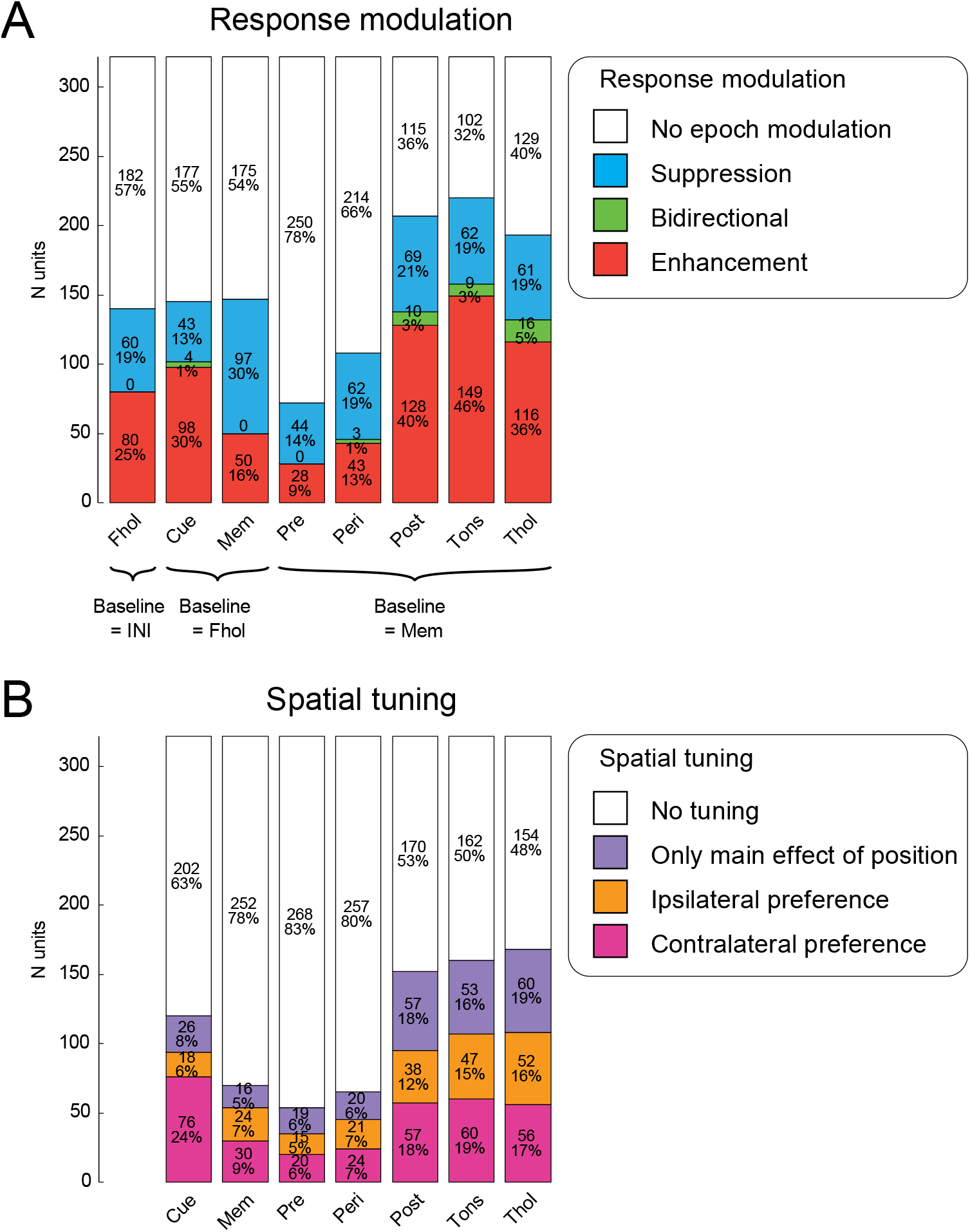
Spatial tuning and response modulation per epoch. Spatial tuning and response modulation of all 322 units recorded during the memory-guided saccade task. The analyzed epochs are fixation hold (Fhol), cue onset (Cue), memory (Mem), pre-saccadic (Pre), Peri-saccadic (Peri), Post-saccadic (Post), target onset (Tons), and target hold (Thol). **A:** Total number of units (and percentages) showing response modulation relative to the respective baseline, for each analyzed epochs: no enhancement nor suppression (white), only suppression (blue), bidirectional - enhancement for one hemifield and suppression for the other (green), only enhancement (red). **B:** Number of units that, in the respective epoch, were not tuned (white), did not prefer either hemifield but showed a main effect of position in a one-way ANOVA (purple), preferred ipsilateral hemifield (orange), preferred contralateral hemifield (magenta). The patterns of response modulation and tuning were largely comparable across the two animals. The most noticeable difference between monkeys was that monkey C showed stronger contralateral cue preference (N=53) compared to ipsilateral cue preference (N=7) than monkey L (N=23 and N=11, respectively). See **Supplementary Figure S1** for the relationship between response modulation and spatial tuning in each epoch.

### Cue and saccade response field properties

To further characterize spatial selectivity in the two epochs with most response modulation, we compared visual and post-saccadic response fields (RFs). We fitted Gaussian response field zones for each unit that showed enhancement or suppression for at least one hemifield in the respective epoch (N=145 for visual and N=207 post-saccadic responses, **Figure 4A,B**), allowing one enhanced and one suppressed zone (**Materials and Methods**). **Figure 4C** shows a summary of RF zones for both epochs. While visual RFs were dominated by contralateral enhancement (N=67, 60% of significant fits), the centers of enhanced post-saccadic response field zones were distributed over both hemifields. Fitted unimodal RFs matching the hemifield-based category (one enhancement zone for units categorized as showing enhancement in at least one hemifield, or one suppression zone for units showing suppression) were more frequently found in visually enhanced units (75 out of 98 ‘en’ units) than in visually suppressed units (25 out of 43 ‘su’ units), indicating that suppression was less topographically consistent (p=0.042; Fisher’s exact test). The visual RFs were less extensive than post-saccadic RFs (mean ± SD 25.7 ± 9.5° and 28.7 ± 10.8° respectively; p=0.018, unpaired t-test).

**Figure 4.**
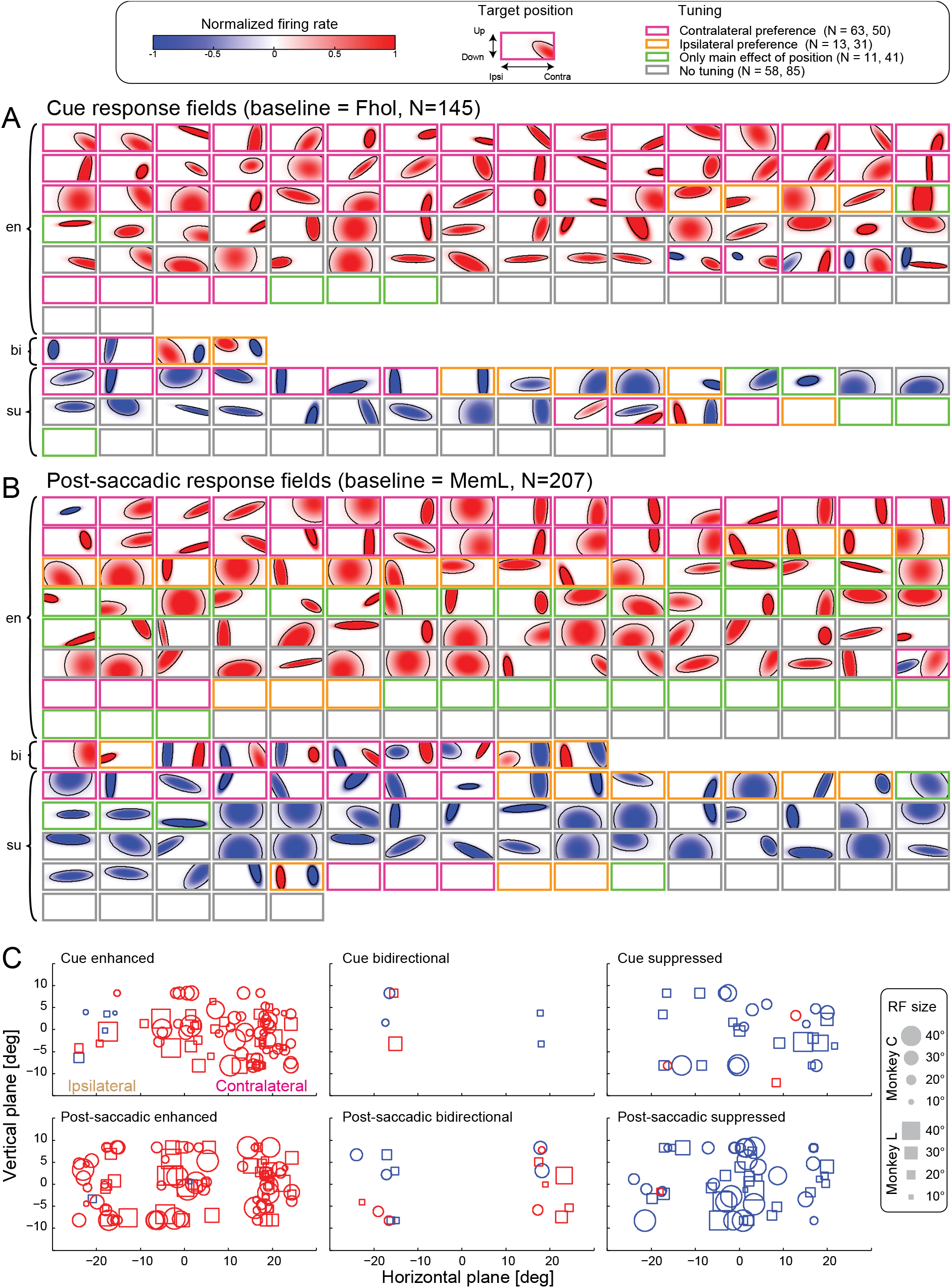
Visual and post-saccadic response fields (RFs). **A:** Visual RFs for units that showed enhancement (en) during cue onset for at least one hemifield, suppression (su), or a bidirectional response (bi – enhancement in one and suppression in the other hemifield). Each subplot represents one unit. Response fields were allowed to have two zones, one enhanced (red) and one suppressed (blue) relative to the respective baseline, see **Materials and Methods**. Only significant zones are plotted; empty subplots indicate no significant explanation of variance by the fitted response field. Firing rates are normalized to the maximum absolute response strength. The color of the frame for each subplot indicates the spatial tuning of the respective unit (significant unpaired t-test on contralateral trials versus ipsilateral trials and/or an ANOVA effect of target position): contralateral preference (magenta), ipsilateral preference (orange), only an ANOVA effect of target position (green), or no spatial tuning (grey). The number of units showing enhancement or suppression in at least one hemifield and the respective tuning (contralateral preference, ipsilateral preference, main effect of ANOVA only, no tuning) is shown in the legend for the cue epoch (left number) and the post-saccadic epoch (right number). Note that units with an ipsilateral suppressed RF zone are expected to show contralateral preference according to our definition of hemifield preference (preferred hemifield is the hemifield with higher firing rate), and vice versa. **B:** Same as A, for post-saccadic RFs. **C:** Summary of all cue (top) and post-saccadic (bottom) response field zones, separately for enhanced, bidirectional and suppressed units. Each symbol represents one zone, circles for monkey C and squares for monkey L. The location and size of the symbols depicts the center and the size of the response field zone, colors indicate enhancement (red) or suppression (blue).

Given that most RFs were large, and some were bilateral with centers near the fovea, we next assessed the relative impact of the direction of a target relative to central fixation vs. its absolute eccentricity. To this end, we performed a direction × absolute eccentricity two-way ANOVA on firing rates in the cue and post-saccadic epoch (**Supplementary Figure S2**). Out of the 113 units showing at least one of the two main effects or an interaction in the cue epoch, most units (N=63) showed only the main effect of direction, 33 units showed some combination of direction and eccentricity effects, and only 17 units showed exclusively the main effect of eccentricity. Likewise, in the post-saccadic epoch out of 175 units with at least one main effect or interaction most units (N=68) showed only the main effect of direction, but a larger proportion showed only the main effect of eccentricity (35 units). This further illustrates that the non-directional eccentricity dependence, while somewhat more frequent in post-saccadic responses compared to visual responses, was a not a common pattern.

To assess potential interactions between hemifields, we asked if the visual stimulus in the opposite hemifield affects the response to the stimulus in the preferred hemifield. Across units with significant cue enhancement, on average there was no difference between response amplitudes in unilateral vs. bilateral trials (**Supplementary Figure S3**). But in line with our expectations, the horizontal RF size (the projection of the fitted RF to the horizontal axis) was larger for the units showing significant bilateral > unilateral response (inter-hemifield summation, 12 units, 17.9±7.0°) than for the units showing unilateral > bilateral response (inter-hemifield suppression; 19 units, 12.6±6.1°, p=0.033, unpaired t-test). Thus, while most units were not affected by the stimulus outside of the preferred hemifield, few units with smaller RFs showed inter-hemifield suppression, and few units with larger RF – summation across the two hemifields.

### Response pattern categorization

To characterize the firing patterns across the sample while taking into account relative predominance of cue and post-saccadic responses (as compared to less frequent pre-saccadic or peri-saccadic responses), we defined three response categories based on the response modulation (enhancement or suppression) during the cue onset and post-saccadic epochs. Visual neurons were defined by statistically-significant enhancement or suppression during cue onset, but no modulation during the post-saccadic epoch, visuomotor neurons were defined by enhancement or suppression in both epochs, and motor neurons by enhancement or suppression in the post-saccadic epoch, but not during cue onset.

**Figure 5A-C** shows two example units for each of these three categories, one example with enhancement during the respective epoch(s) on the left, and suppression on the right. The diversity of the response patterns is further illustrated by example units which showed enhancement or suppression during the memory epoch (**Figure 5D**) and units with spatial preference during the target hold epoch (**Figure 5E**). The localization of units categorized as visual (N=50), motor (N=112), or visuomotor (N=95) neurons revealed no systematic distribution of the recording sites within the medial dorsal pulvinar (**Figure 6A**). To further confirm the plausibility of this categorization, we computed a visuomotor index (VMI, ranging from −1 for purely visual to 1 for purely motor response) for each unit independently for the ipsilateral and the contralateral hemifield (see **Materials and Methods**) and tested if the mean VMIs in each group were significantly different from zero. As expected, VMIs of visual units were significantly negative and VMIs of motor units were significantly positive, see **Figure 6B** (p<0.001 for all four tests, one sample t-test). Interestingly, VMIs of visuomotor units were significantly positive for the ipsilateral hemifield (p<0.01), while for the contralateral hemifield VMIs of visuomotor units were not different from zero (p=0.46). This indicates stronger motor responses for ipsilateral saccades as compared to visual responses to ipsilateral stimuli in visuomotor units. The categories however overlapped and there were no separate modes in VMI distributions, indicating a continuum of visuomotor properties. We retain these categories for subsequent analyses as providing a tractable approach to the heterogeneity of response patterns, but do not imply separable groups.

**Figure 5.**
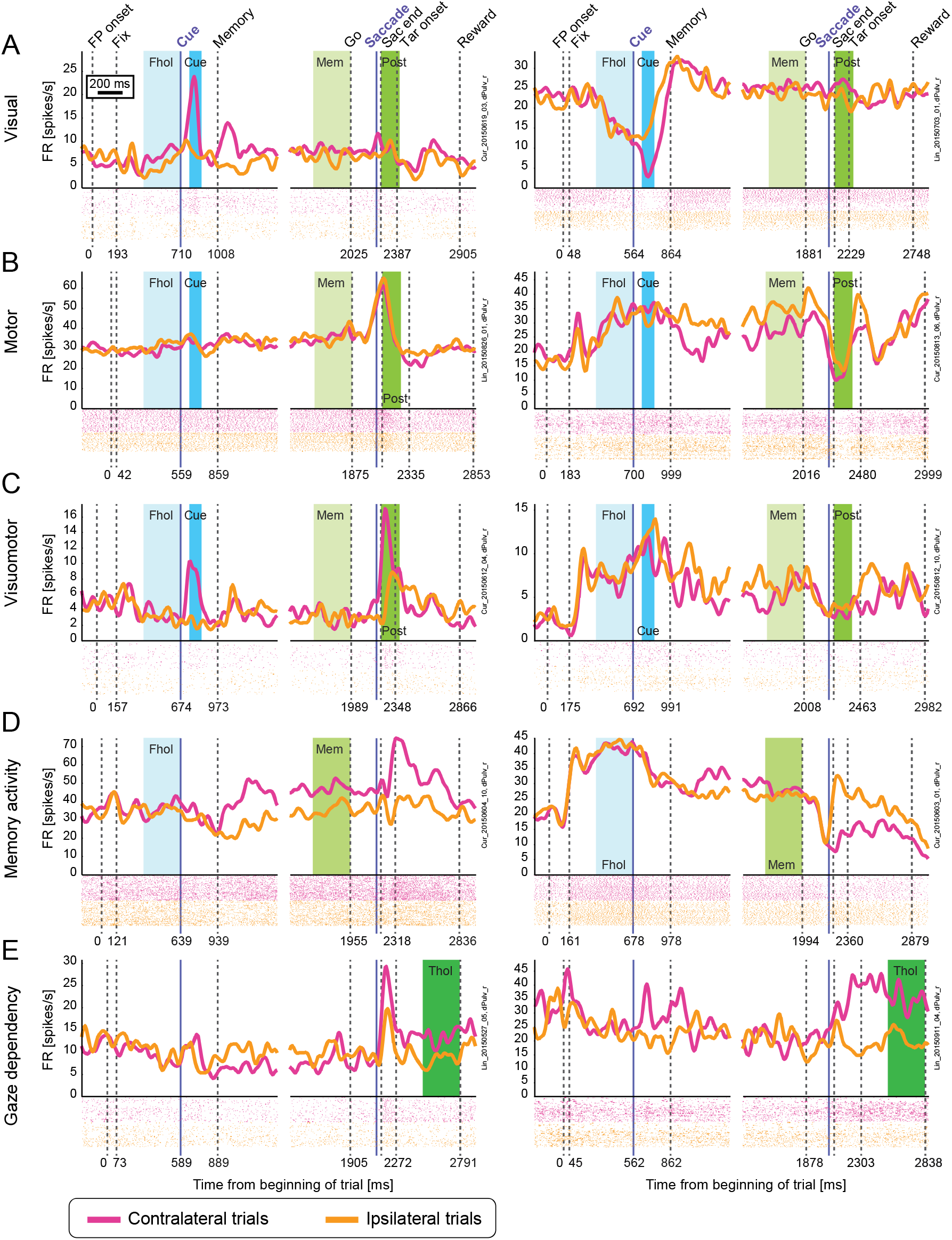
Diversity of task-related firing modulations: example units. Raster plots and spike density functions for contralateral (magenta) and ipsilateral (orange) trials. Vertical lines indicate the average onset of events: fixation point onset (FP onset), the monkey acquiring fixation (Fix), the cue onset, the cue offset and beginning of the memory period, the offset of the central fixation point (Go), the saccade onset, the monkey acquiring the invisible target location (Sac end), the onset of the visible peripheral target, and the end of the trial (Reward). Discontinuous traces indicate two different alignments to cue onset and saccade onset (purple lines). Colored areas mark relevant analyzed epochs: fixation hold (Fix), cue onset (Cue), memory (Mem), post-saccadic (Post), and target hold (Thol). Examples for five different categories are shown. **A:** Visual units (enhancement or suppression in Cue relative to Fhol; **B:** Motor units (enhancement or suppression in Post relative to Mem); **C:** Visuomotor units (enhancement or suppression in both Cue and Post); **D:** Units with memory activity (left: hemifield preference in Mem, right: suppression in Mem compared to Fix); **E:** Gaze dependent activity (hemifield preference in Thol, in the period starting 300-400 ms after the end of the saccade, 200 ms after target onset).

**Figure 6.**
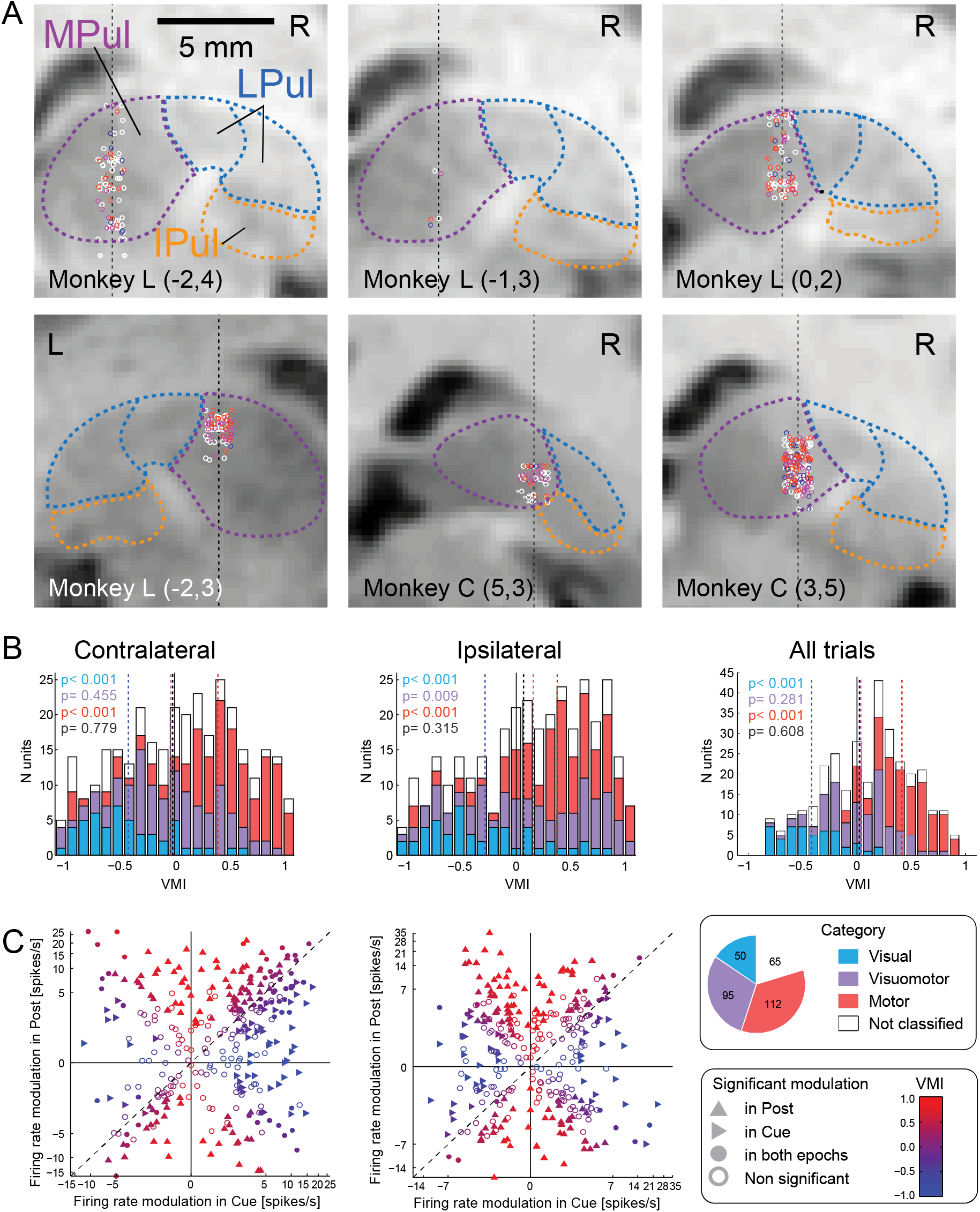
Visuomotor categories. **A:** Localization of recorded units in chamber-normal coronal sections in each monkey (L and C, labels on the bottom) and specific grid locations (x,y in parentheses). Locations were jittered along the horizontal axis for better visualization. The black dotted lines indicate the electrode tracks and mark the actual horizontal location of recorded neurons. Colors indicate the category of the unit; blue for visual, magenta for visuomotor and red for motor units. Units that did not fall within any of these categories are shown as white circles. Pulvinar nuclei outlines (MPul/LPul/IPul – medial/lateral/inferior pulvinar) were adapted from the NeuroMaps atlas (Rohlfing et al. 2012), exported via Scalable Brain Atlas, https://scalablebrainatlas.incf.org/macaque/DB09, https://scalablebrainatlas.incf.org/services/rgbslice.php, (Bakker et al. 2015), and LPul was further subdivided to dorsal (PLdm) and ventral (PLvl) parts. **B:** Histograms of VMIs computed for contralateral trials (left), ipsilateral trials (middle), or all trials (right). Colors denote the category. P-values indicate results of one sample t-tests against zero. **C:** Scatter plots comparing the firing rate modulation in Cue and Post-saccadic epochs, for contralateral trials (left) and ipsilateral trials (right). Colors indicate the respective visuomotor index (legend inset on the right). For better visualization, firing rate modulation was plotted on a logarithmic scale. The pie plot on the right shows the total number of units for each of the categories.

To further test if specific sub-groups would be evident across the entire population, we plotted post-saccadic modulation versus cue modulation across the entire population (**Figure 6C**). Besides a noticeable cluster of units in the upper right corner showing similar enhancement during the cue onset epoch and the post-saccadic epoch in contralateral trials, the units were distributed fairly uniformly, further indicating a continuum of responses.

Population PSTHs for the three categories show enhancement for contralateral cues in visual and visuomotor units, and post-saccadic enhancement with only weak spatial preference in visuomotor and motor units (**Figure 7A**, top). The visual neurons show only a very small bump after a saccade but respond later to a target onset, motor neurons respond to the saccade, and visuomotor neurons to both events. A more detailed picture can be gained by further separating these three categories based on enhancement or suppression either in the cue onset or the post-saccadic epochs, and analyzing the preferred target position for each unit (the position that exhibited the strongest response modulation in the respective epoch, **Figure 7A**, bottom). Two observations can be made. First, visuomotor units showing suppression in the cue epoch relative to fixation hold are typically also suppressed throughout the entire trial, including memory delay (although the visual conditions are identical in the fixation hold and memory periods). This highlights a group of neurons with enhanced fixation hold responses. Second, the net enhancement of visuomotor and motor units in the post-saccadic epoch is a result of unequal proportions of enhanced and suppressed units (N=62 and N=29, 65% and 31% of visuomotor units; N=66 and N=40, 59% and 36% of motor units).

**Figure 7.**
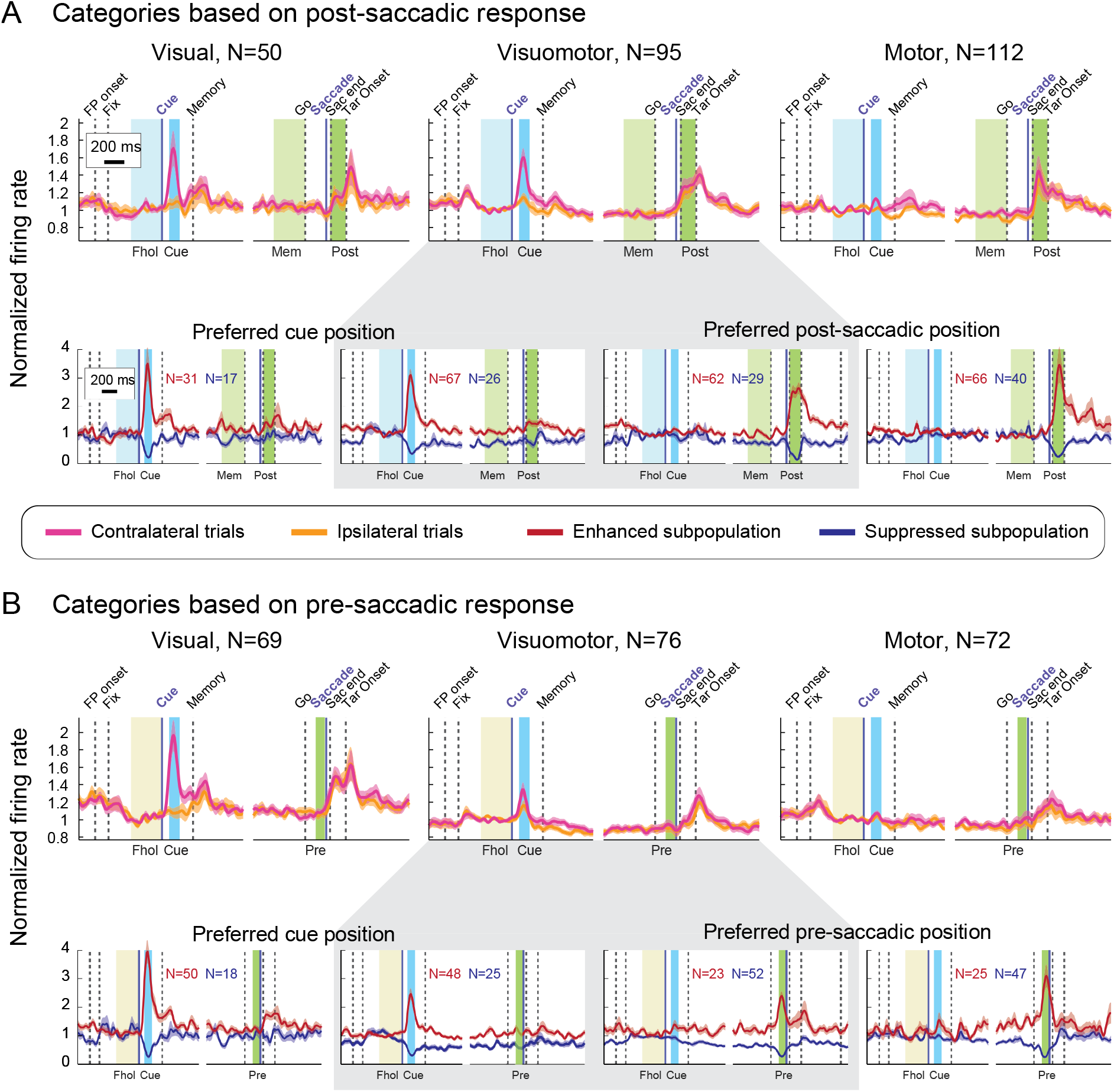
Population responses of visual, visuomotor, and motor units. Vertical lines mark events, and colored areas relevant epochs for categorization. **A:** Population response of the three categories: visual (enhancement or suppression in Cue relative to Fhol, but not in Post), motor (enhancement or suppression in Post relative to Mem, but not in Cue) and visuomotor units (enhancement or suppression in both epochs). Top: population responses for contralateral (magenta) and ipsilateral (orange) trials. Bottom: subpopulations showing enhancement (red) or suppression (blue) either during Cue or Post-saccadic epochs. For each unit, only the response for the most modulated target position (location with the largest firing rate difference from baseline) in the respective epoch was taken (note that few units showing enhancement for one and suppression for another location were excluded in those plots). **B:** same as A, for the categorization based on pre-saccadic response: visual (enhancement or suppression in Cue relative to Fhol, but not in Pre), motor (enhancement or suppression in Pre relative to Fhol, but not in Cue) and visuomotor units (enhancement or suppression in both epochs). Note that the difference in the frequency of pre-saccadic enhancement in the **Figure 3A** (9%) and here (23+25 units, 15%) stems from different baseline periods for assessing the enhancement (the memory period in **Figure 3A** and the fixation hold here).

To isolate most common patterns in these three categories, and to address the contribution of units with different spatial preferences, we defined subpopulations of each of the categories based on enhancement or suppression and spatial preference either in the cue onset epoch or the post-saccadic epoch (**Table 1**, see also **Supplementary Figure S4**). These numbers show that the weak overall post-saccadic spatial preference was due to a larger proportion of contralaterally-tuned than ipsilaterally-tuned units, on top of a large faction of units without tuning. Post-saccadic hemifield preference was more common in visuomotor units than in motor units (48% vs. 31%, p=0.015; Fisher’s exact test), suggesting that the former have a larger role in spatially-specific processing.

**Table 1.** Visual, visuomotor, and motor subpopulations. Subpopulations were defined by response modulation and spatial tuning either in the cue onset epoch (for visual and visuomotor units) or in the post-saccadic epoch (for visuomotor and motor units). Numbers indicate the total number for each subpopulation and the percentage of the respective “parent” category. The red/blue/green colors correspond to the pattern of response modulation (enhancement, suppression, bidirectional) in **Figure 3A**. See **Supplementary Figure S4** for visualization of these response patterns.

| Separation by response modulation and tuning in cue onset epoch |  |  |  |
| --- | --- | --- | --- |
|  | Visual (N=50) | Visuomotor (N=95) | Motor (N=112) |
| Enhancement, no tuning | 17 (34%) | 26 (27%) | - |
| Enhancement, contralateral preference | 14 (28%) | 37 (39%) | - |
| Enhancement, ipsilateral preference | 0 (0%) | 4 (4%) | - |
| Suppression, no tuning | 8 (16%) | 18 (19%) | - |
| Suppression, contralateral preference | 6 (12%) | 4 (4%) | - |
| Suppression, ipsilateral preference | 3 (6%) | 4 (4%) | - |
| Contra enhancement and ipsi suppression | 2 (4%) | 0 (0%) | - |
| Contra suppression and ipsi enhancement | 0 (0%) | 2 (2%) | - |
| Separation by response modulation and tuning in post-saccadic epoch |  |  |  |
|  | Visual (N=50) | Visuomotor (N=95) | Motor (N=112) |
| Enhancement, no tuning | - | 32 (34%) | 46 (41%) |
| Enhancement, contralateral preference | - | 19 (20%) | 13 (12%) |
| Enhancement, ipsilateral preference | - | 11 (12%) | 7 (6%) |
| Suppression, no tuning | - | 17 (18%) | 31 (28%) |
| Suppression, contralateral preference | - | 3 (3%) | 8 (7%) |
| Suppression, ipsilateral preference | - | 9 (9%) | 1 (1%) |
| Contra enhancement and ipsi suppression | - | 3 (3%) | 4 (4%) |
| Contra suppression and ipsi enhancement | - | 1 (1%) | 2 (2%) |

Since we were also interested in characterizing pre-saccadic responses, **Figure 7B** illustrates the same analysis based on the alternative motor-related categorization that considered response modulation in the *pre*-saccadic epoch relative to fixation hold (**Materials and Methods**). Importantly, the post-saccadic enhancement is still evident in the population responses of all three categories, while enhancement and suppression as well as spatial preference in the pre-saccadic epoch balances out completely across the sample (**Figure 7B**, top), due to opposing contributions of enhanced and suppressed units (**Figure 7B**, bottom). This further supports a classification based on post-saccadic responses.

### Comparison of memory-guided and visually-guided saccade tasks

To evaluate how the presence of a visible saccade target and the immediate response context might affect pre-saccadic activity as compared to memory-guided saccade, in a subset of sessions we also recorded the same units while monkeys were performing a visually-guided saccade task. In the visually-guided task, monkeys had to perform a saccade immediately after the onset of the target(s) (**Materials and Methods**).

We compared the response modulation in the target onset/pre-saccadic epoch in visually-guided task against two partly corresponding epochs in the memory-guided task: ‘cue onset 2’ that encompassed most visual responses, 40-140 ms after the cue and ‘pre-saccadic 2’ that encompassed motor preparation signals, 130-30 ms before the memory saccade onset. These epochs differ slightly from the original cue (70 ms to 170 ms after cue onset) and pre-saccadic (100 to 10 ms before saccade onset) epochs, and were chosen to be comparable to the combined target onset/pre-saccadic epoch 40-140 ms after the target onset (‘target-pre-sac’) in the visually-guided task – where this interval was chosen to not overlap with the saccade onset. Response modulation in each epoch was calculated by subtracting firing rates in fixation hold (Fhol) to allow for direct comparison between the two tasks, in all units recorded in both tasks (N=174).

Response modulations were correlated across units in the two tasks, when correlating ‘target-pre-sac’ response to either visual (‘cue onset 2’) or pre-saccadic responses (‘pre-saccadic 2’) (p<0.001 for ‘cue onset 2’ for both hemifields, p=0.063 for ‘pre-saccadic 2’ ipsilateral, p=0.002 for ‘pre-saccadic 2’ contralateral, Pearson’s correlation). This indicated that both visual and motor preparatory signals contribute to response modulations in the visually-guided task. To assess if the response modulation in the visually-guided task can be explained by the sum of visual and motor preparation signals, we compared activity in the ‘target-pre-sac’ to the sum of ‘cue onset 2’ and ‘pre-saccadic 2’ responses. The ‘target-pre-sac’ response modulation was stronger than the sum of separate epochs, suggesting supralinear summation of visual and premotor activity (p=0.002 for contralateral trials, p<0.001 for ipsilateral trials; paired t-tests). This was particularly true for units showing enhancement in the visually-guided task (contralateral trials: N=49, p<0.001; ipsilateral trials: N=19, p=0.007; paired t-tests). Thus, the immediate response context of the visually-guided task enhanced pre-saccadic activity.

### Delay period and pre-/peri-saccadic activity

Due to the strong bidirectional connectivity between dorsal pulvinar and posterior parietal cortex (PPC), in particular area LIP (Hardy and Lynch 1992; Gutierrez et al. 2000) as well as the suggested role of pulvinar in guiding goal-directed actions (Grieve et al. 2000; Wilke, Turchi, et al. 2010), we predicted that some dPul neurons show delay period and pre-/peri-saccadic activity – a hallmark of canonical PPC responses (Barash et al. 1991a; Premereur et al. 2011). As reported previously (Dominguez-Vargas et al. 2017), when looking at population averages for units that showed either enhancement or suppression in the peri-saccadic epoch as compared to fixation hold, it seems that a considerable subset indeed showed activity ramping up (N=42) or ramping down (N=94), reaching the respective maximum or minimum during saccade execution (**Figure 8A,B**). Normalizing these two subsets using division by average firing rate in fixation hold rather than subtracting baseline inter-trial-interval activity strengthened that impression (**Figure 8C,D**). However, here we demonstrate that the continuously ramping up delay period activity culminating at the saccade is largely a result of combining different subpopulations. First, subpopulations showing peri-saccadic enhancement (N=43) or suppression (N=62) relative to memory did not exhibit ramping or persistent delay period activity (**Figure 8E,F**). Second, subpopulations showing enhancement or suppression in memory relative to fixation hold (N=50 and N=97 respectively) did not exhibit pronounced peri-saccadic responses (**Figure 8G,H**). In fact, very little overlap between these subpopulations was found: only 4 units showed significant enhancement in memory relative to fixation hold and in peri-saccadic epoch relative to memory, and 18 units showed suppression in both epochs. Besides these few consistently ramping up/down units, subpopulations shown in **Figure 8A,C** and **B**,**D** combine units that showed enhancement or suppression during memory (relative to fixation) with units that showed enhancement or suppression during the peri-saccadic epoch (relative to memory). Interestingly, the enhanced memory delay firing stayed at the same level throughout the entire period (**Figure 8G**), whereas activity was decreasing gradually in the subpopulation showing suppression in the memory epoch (**Figure 8H**).

**Figure 8.**
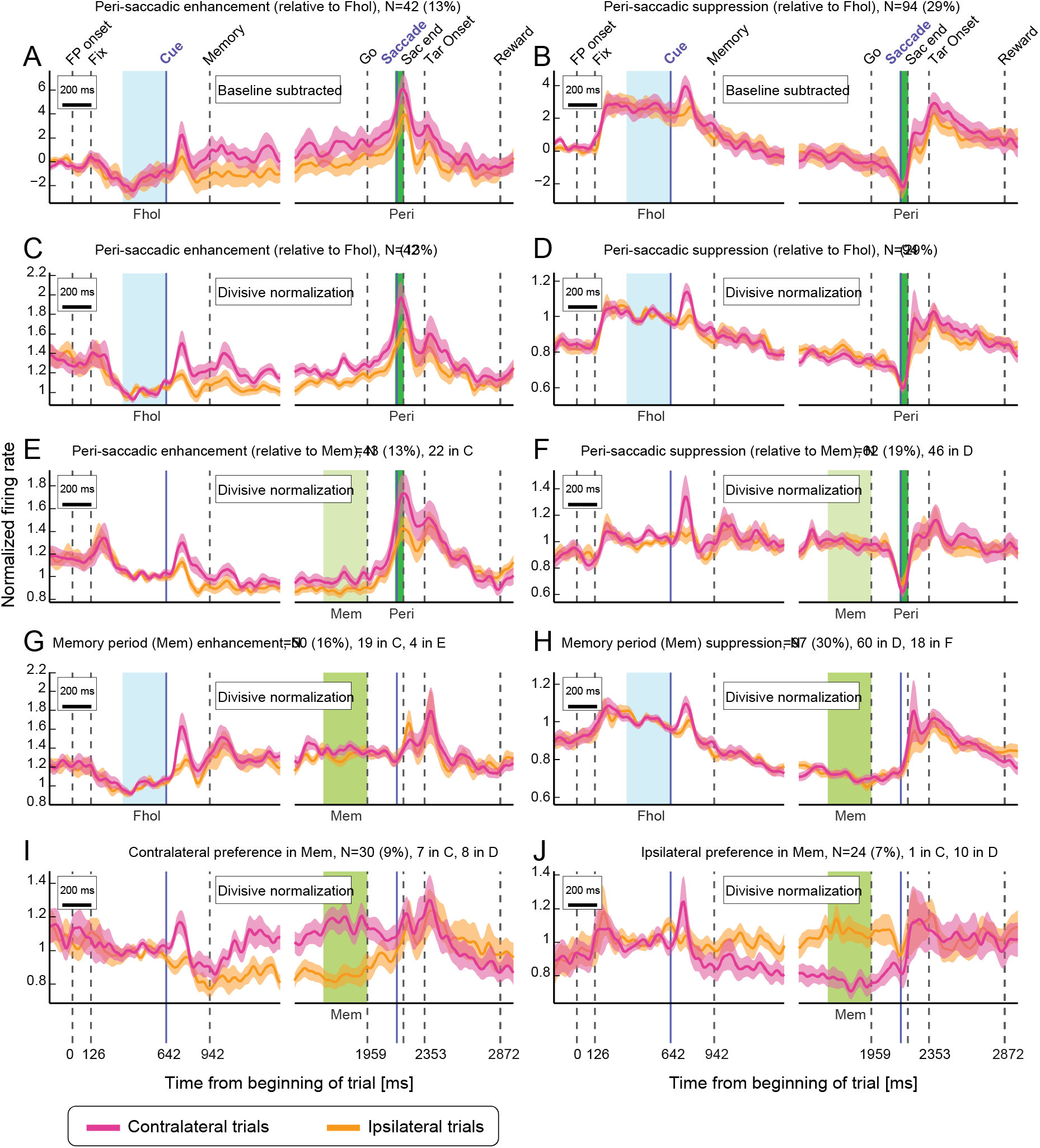
Delay-period activity. Population responses for contralateral (magenta) and ipsilateral (orange) trials. Vertical lines mark events, and colored areas relevant epochs for population definition. **A:** Average of units showing peri-saccadic enhancement relative to fixation hold epoch (Fhol), normalized by subtracting the average firing rate in inter-trial-interval epoch. **B:** Same as A, for units showing peri-saccadic suppression. **C:** Average of units showing peri-saccadic enhancement relative to Fhol, normalized by dividing the spike density functions by the average firing rate in Fhol. **D:** Same as C for units showing peri-saccadic suppression. **E:** Average of units showing peri-saccadic enhancement relative to the memory epoch (Mem). **F:** Same as E for units showing peri-saccadic suppression. **G:** Average of units showing enhancement in Mem relative to Fhol. **H:** Same as G for units showing suppression. **I:** Average of units showing contralateral preference in Mem. **J:** Same as I for units showing ipsilateral preference. Additional text on top of E-J panels shows number of units in the current panel overlapping with other indicated panels.

In studies of parietal cortex, neurons showing enhanced firing to visual cues are often selected online or offline for further analyses of delay period activity (Colby et al. 1996; Ipata et al. 2006; Chen et al. 2016); see also a recent study in the pulvinar (Fiebelkorn et al. 2019). Our sample was acquired without taking into account any response properties online; across the subset of 98 units that exhibited enhanced cue response in the offline analysis, the curves for the preferred and nonpreferred target positions largely overlapped during the delay. Likewise, across the entire sample the curves for contralateral and ipsilateral positions largely overlapped in the late memory delay (**Supplementary Figure S5**). **Figure 8I,J** shows specific subpopulations that were significantly tuned for contralateral or ipsilateral space in the memory delay. The contralaterally- and ipsilaterally-tuned units differed in terms of response pattern: the contralaterally-tuned group showed a congruent contralateral cue tuning (but no pre-/peri-saccadic enhancement), while the ipsilaterally-tuned group showed incongruent contralateral cue tuning as well as peri-saccadic suppression in ipsilateral trials. Furthermore, out of 50 units with memory period enhancement, most tuned units had contralateral preference (11 contralateral, 2 ipsilateral), unlike more balanced tuning in 97 units with memory period suppression (14 contralateral, 12 ipsilateral; **Supplementary Figure S1**).

These findings have two important implications. Firstly, only a small minority of units showed “classical” spatially-tuned cue-delay-saccade enhancement, and the units that were contralaterally-tuned in the cue and the memory period exhibited sustained delay activity but no ramping towards the saccade, constraining the role of individual dPul neurons in prospective motor planning. Secondly, more units showed delay period suppression and *ramping down* than enhancement, and most of these units showed cue suppression or no cue response (76 out of 97 units with memory delay suppression), suggesting putative inhibitory projections from a subset of pulvinar units to the frontoparietal circuitry, or conversely a cortically-driven inhibition on the pulvinar.

### Spatial choice selectivity

So far we mostly considered single cue instructed trials (except in the section on the RF properties). In another half of the trials, monkeys were presented with two cues equidistantly from the fixation point, and were free to select either of the two positions, for the same reward. There were 6 possible pairs of targets, and instructed and choice trials were randomly interleaved, discouraging monkeys to make selection before the cues were presented. It has been shown that in such conditions dPul inactivation biases the choice towards ipsilesional hemifield (Wilke et al. 2013).

Target selection in choice trials was fairly balanced between the two hemifields in both monkeys (monkey C: 60/40%; monkey L: 48/52%) and reaction times in choice trials were very similar to instructed trials. If anything, monkeys were slightly faster in choice trials (**Table 2**). The lack of additional delay in choice RTs suggests that monkeys made choices before the end of delay period.

**Table 2.** Saccade reaction times (RT) and spatial target selection. P values are from paired t-tests comparing session means.

|  | Hemifield | Instructed<br>RT | Choice<br>RT | p value<br>(instr. vs. choice RT) | % selected | p value<br>(selection left vs. right) |
| --- | --- | --- | --- | --- | --- | --- |
| Monkey C | Left | 207 | 204 | 0.017 | 62 ± 3 | <0.001 |
|  | Right | 217 | 209 | <0.001 | 38 ± 3 |  |
| Monkey L | Left | 193 | 195 | 0.302 | 48 ± 5 | 0.72 |
|  | Right | 189 | 186 | 0.002 | 52 ± 5 |  |

To assess if spatial preference in dPul neurons during the delay period is linked to space-specific motor preparation and could therefore reflect the upcoming saccade decision, we analyzed choice trials in the subpopulations showing hemifield preference in instructed trials. On average, there was no hemifield preference in the delay period activity in choice trials, and the choice trial traces were situated between preferred and opposite hemifield instructed trials (**Figure 9A**). This might indicate that in these units the spatially-selective delay activity in instructed trials is not directly linked to prospective target selection and motor preparation, but to retrospective maintenance of information about the visual stimuli presented earlier in the trial, or attentional allocation.

**Figure 9.**
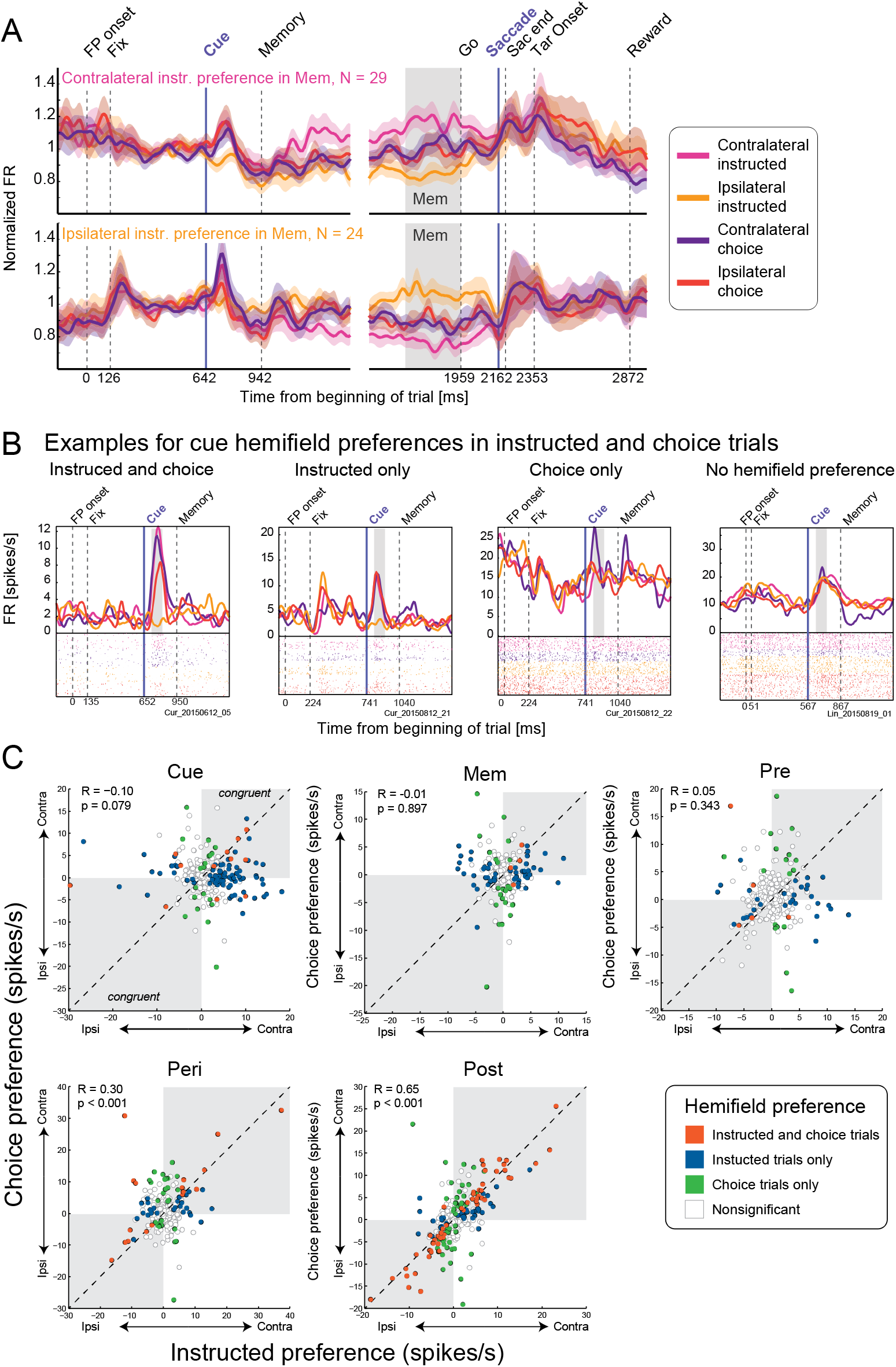
Choice selectivity. **A:** Population averages for 29 units showing either contralateral preference (top) or ipsilateral preference (bottom) in the memory (Mem) epoch, for instructed contralateral (magenta) & ipsilateral (orange), choice contralateral (purple) & ipsilateral (dark red) trials. One unit was excluded because it was recorded for <4 choice trials to one hemifield. **B:** Example units with different types of cue responses, from left to right: Contralateral preference in both instructed and choice (CI_instr_ = 0.84, CI_choice_ = 0.23), contralateral preference only in instructed (CI_instr_ = 0.52, CI_choice_ = −0.01), contralateral preference only in choice (CI_instr_ = 0.09, CI_choice_ = 0.18), no hemifield preference (CI_instr_ = −0.05, CI_choice_ = 0.05). **C:** Scatter plots comparing firing rate hemifield preference (FR_contra_ – FR_ipsi_) in instructed and choice trials, for six epochs: cue onset (Cue), memory (Mem), pre-saccadic (Pre), peri-saccadic (Pre) and post-saccadic (Post). R and p values are for Pearson’s correlations between hemifield preference in instructed and choice trials; congruent tuning quadrants are in gray. Five units were excluded from this analysis because <4 choice trials for one hemifield were recorded (N=317).

Some units however showed a significant difference in firing rates prior to selection of the contralateral or ipsilateral target in choice trials, well before the saccade. The pertinent question is whether the spatial selectivity in choice trials is mostly congruent to the spatial selectivity in instructed trials, as would be expected from neurons participating in motor decision and prospective planning (Coe et al. 2002; Watanabe and Funahashi 2007; Klaes et al. 2011; Pastor-Bernier and Cisek 2011; Suriya-Arunroj and Gail 2019). **Figure 9B** shows four example units with different types of cue tuning: i) with congruent hemifield preference in instructed and choice trials, ii) only in instructed trials, iii) only in choice trials, iv) no hemifield preference in either instructed and choice trials. To evaluate if pulvinar neurons encode the upcoming choice across the population, we compared hemifield preference between instructed and choice trials in each unit in five epochs (cue onset, memory, pre-saccadic, peri-saccadic, and post-saccadic epoch). Before the saccade (cue, memory, pre-saccadic epochs) there was no systematic congruency between tuning in instructed and choice trials (**Figure 9C**). Only few units showed a significant *congruent* effect of the (selected) hemifield in both trial types (cue: 8 units, memory: 3 units, pre-saccadic: 2 units), and across all units, hemifield preferences (firing rate difference contra – ipsi) were not correlated (cue: p=0.079, memory: p=0.897, pre-saccadic: p=0.343, Pearson’s correlation; the same result holds true for only those units that were tuned in the instructed trials). Furthermore, the strength of cue spatial preference (defined as the absolute of the Contralaterality Index, see **Materials and Methods**) was higher in instructed than in choice trials (instructed minus choice index difference 0.04 for all units, p<0.001, Wilcoxon’s signed rank test, and 0.13 for units tuned in instructed and/or choice trials). As a control analysis, hemifield preferences were highly correlated between instructed and choice trials in peri- and post-saccadic epochs (p<0.001, Pearson’s correlation), as expected given that the movements to the same target were similar in instructed and choice trials.

Because the choice selectivity could potentially be “washed out” by including multi-units, we also performed the same analysis on single units only (N=245, see **Supplementary Figure S6**), and found similar results, with exception of a very weak but significant (p=0.002) correlation in the pre-saccadic epoch. Furthermore, the same analysis on ‘target-pre-sac’ epoch in visually-guided saccades also showed a significant correlation (p=0.006), despite the fact that the activity in this epoch represents not only the upcoming saccade preparation but also the visual input that is the same regardless of contralateral or ipsilateral choice. These findings suggest that the upcoming spatial decision and movement planning were not encoded in the dPul until shortly before the saccade. Instead, the spatially-selective delay period activity during the instructed trials can be largely attributed to retrospective memory of visual stimulation or attentional allocation.

## Discussion

The goal of this study was to characterize visual, delay and motor properties and choice selectivity of dorsal pulvinar neurons, similar to what has been done extensively for other thalamic nuclei and frontoparietal cortical regions (Barash et al. 1991a, 1991b; Funahashi et al. 1991; Chafee and Goldman-Rakic 1998; Constantinidis et al. 2001; Wyder et al. 2003; Watanabe et al. 2006; Tanaka and Kunimatsu 2011). We utilized a classical memory-guided saccade task to dissociate sensory encoding, intervening delay and motor execution phases, and found a considerable diversity of response patterns. Our results demonstrate that most dorsal pulvinar neurons exhibit either enhancement or suppression around or after saccades. Saccade-related responses were mostly post-saccadic, although there were also a substantial number of neurons showing pre- and peri-saccadic modulation. Only a small fraction of neurons responded exclusively to visual stimulation, further supporting the notion that the dorsal pulvinar is involved in oculomotor processing. Delay period activity was mainly characterized by a gradual suppression of firing relative to the initial fixation period, with only a small subset showing classical sustained or ramping up enhancement of activity that is frequently observed in frontoparietal areas such as LIP and FEF. Surprisingly, on the population level no spatial choice selectivity for the upcoming saccade was found, neither in the cue/delay nor in pre-saccadic activity, and individual units rarely showed a consistent spatial tuning in single target instructed and two-target choice trials.

### Anatomical considerations

When comparing results presented here with the few previous studies on visual and oculomotor motor properties of the pulvinar, it is important to note that most studies in the dorsal pulvinar recorded from the dorsomedial division of the *lateral* pulvinar. This division is denoted as PLdm (Kaas and Lyon 2007; Bridge et al. 2015; Kagan et al. 2021), or as Pdm in original electrophysiological studies by Robinson, Petersen and colleagues, e.g. (Petersen et al. 1985, 1987; Robinson et al. 1986; Bender and Youakim 2001). The notable exception is a study that covered both lateral and medial parts of the dorsal pulvinar, predominantly at the border between the two (Benevento and Port 1995). Our recording sites were largely within the medial pulvinar (MPul) or around the border between the medial and the lateral pulvinar (see also (Dominguez-Vargas et al. 2017; Schneider et al. 2019)). We did not observe any clear patterns along the dorsal/lateral – ventral/medial axis in the features we tested, and such patterns have also not been reported in the previous work. However, given rostral-caudal and lateral-medial gradients of dPul anatomical and functional connectivity (Baleydier and Morel 1992; Baizer et al. 1993; Gutierrez et al. 2000; Arcaro et al. 2018; Froesel et al. 2021), more extensive electrophysiological and combined perturbation-imaging mapping (Klink et al. 2021) will be required to further assess the functional differences between subdivisions of the dorsal pulvinar.

### Visual and prevalent motor responses

In the memory saccade task, we found diverse patterns of visual and post-saccadic activations along a visuomotor continuum, and no clear organization of these patterns within the dorsal pulvinar. Notably, purely visual responses were less frequent than pure motor responses or combined visual and motor responses (15% visual compared to 30% visuomotor and 35% motor units), supporting an involvement of dorsal pulvinar in distributed neuronal processing related to motor commands and consequences of oculomotor behavior, beyond purely visuospatial functions (Grieve et al. 2000; Guillery and Sherman 2002; Saalmann and Kastner 2015).

Visual responses to a small briefly flashed saccade cue were mainly marked by transient increases of spiking activity, with a clear contralateral preference. This expected presence of cue-related activity contrasts with the previous report in dPul, where responses to visual stimuli were reported only when the stimulus served as a target of an immediate visually-guided saccade but not in memory saccades (Benevento and Port 1995). We indeed found that the combined visual and pre-saccadic activity during visually-guided saccades was stronger than the sum of cue- and pre-saccadic activity during memory saccades – in agreement with the idea that immediate sensorimotor contingency of a stimulus is an important determinant of dPul activity. But robust cue responses during memory-guided saccades were found in 45% of our sample. Indeed, the proportion of units, the size of the receptive fields, the range of latencies, and the contralateral tuning of cue responses in dPul are largely consistent with the data from MD, the central thalamus and frontoparietal cortex. This suggests that a pathway via the dorsal pulvinar similar to well-identified cortico - basal ganglia - oculomotor thalamo-cortical loops, a SC – pulvinar, or a direct cortico-pulvinar connectivity, underlie the visual activity in dPul.

The saccade-related responses in dPul show a mix of cortical and other thalamic nuclei properties. In contrast to mainly pre-saccadic activity in MD (Watanabe and Funahashi 2004a; Cavanaugh et al. 2020) and in the central oculomotor thalamus, where saccade-related responses are characterized by equally common space-specific pre-saccadic (47%) and post-saccadic (53%) activity (Wyder et al. 2003), saccade-related responses in dPul are mainly post-saccadic (64% of all units compared to 22% with pre-saccadic activity). But similarly to the central oculomotor thalamus, dPul shows twice as much post-saccadic enhancement as suppression (40% and 21% of all units) relative to the preceding memory delay period. Likewise, a large fraction of spatially-tuned post-saccadic responses, but with only a weak contralateral spatial preference on a population level, with extensive RFs that are often ipsilateral or extend to the ipsilateral side and include the foveal region, resemble the saccadic RFs in the lateral MD and central oculomotor thalamus (Wyder et al. 2003; Cavanaugh et al. 2020).

Both directionally-selective and non-directional (or “omni-directional”) post-saccadic enhancement responses are common in parietal cortex, in LIP and particularly in area 7a (Barash et al. 1991a, 1991b; Genovesio et al. 2007; Zhou, Liu, et al. 2016), prefrontal cortex (Funahashi 2014) and the thalamus (Wyder et al. 2003; Tanaka and Kunimatsu 2011). A number of functions related to feedback and internal monitoring have been attributed to these responses. The non-space-specific post-saccadic enhancement could be used for timing, to signal that the saccade has ended and processing of visual inputs can be re-established (Schlag and Schlag-Rey 1984; Tanaka and Kunimatsu 2011; Boi et al. 2017; Kagan and Burr 2017; Zhou et al. 2018). In the context of the memory saccade task, post-saccadic enhancement might also reflect the anticipation of a change in visual input such as the confirmation target onset, since it appears soon after the end of the saccade with a predictable timing. Potentially, deviations of the post-saccadic eye position from the intended target location, i.e. error signals, could also be encoded in post-saccadic responses, as has been found in LIP (Zhou, Liu, et al. 2016; Munuera and Duhamel 2020), but the small saccade endpoint dispersion due to fairly stereotypical movements in our experiment did not allow to systematically check for this possibility. Finally, the spatially-tuned post-saccadic responses could also signal the new gaze position, alone or in a combination with previous eye movement vector or pre-saccadic position, and utilized for internal monitoring during action sequences (Genovesio et al. 2007; Tanaka 2007; Schneider et al. 2019).

The second most frequent saccade-related response in dPul was a transient, typically non-space-specific pre- and peri-saccadic suppression (pre: 14%, 71% out of those units not space-specific; peri: 19%, 68% out of those units not space-specific). Some of these neurons also showed a burst after the saccade – similar to so called ‘pause-rebound’ response found in central thalamus (Tanaka and Kunimatsu 2011). The pre/peri-saccadic suppression could reflect extraretinal signals such as corollary discharge intended to prevent self-induced stimulus motion, e.g. by suppressing visual information transfer to visual frontoparietal areas during the saccade (Sommer and Wurtz 2004; Bellebaum et al. 2005; Kagan et al. 2008; Ibbotson and Krekelberg 2011; Wurtz, Joiner, et al. 2011; Wurtz, McAlonan, et al. 2011).

A smaller subset of units exhibited transient pre-saccadic (7%) and peri-saccadic enhancement (13%), with an overall weak preference for the contralateral hemifield. The activity in these units resembles saccade-related responses in oculomotor thalamus, FEF and LIP, although pre/peri-saccadic enhancement is much more prevalent in these regions (Barash et al. 1991a; Wyder et al. 2003; Watanabe et al. 2006). The pre/peri-saccadic response strength in dPul might depend on the behavioral demands and decision urgency, as found in the oculomotor thalamus (Costello et al. 2015). Since we used a fixed memory delay and a relatively generous saccade RT cutoff (500 ms), we surmise that the urgency in our task was rather low, which might explain the low prevalence of pre/peri-saccadic activity in this dataset.

### Delay period activity and choice selectivity

Given the widespread reciprocal connectivity between dorsal pulvinar and the frontoparietal network (Bos and Benevento 1975; Schmahmann and Pandya 1990; Hardy and Lynch 1992), and the fact that dorsal pulvinar perturbation reliably biases saccade target selection during visually-guided and memory-guided saccades (Wilke, Turchi, et al. 2010; Wilke et al. 2013; Dominguez-Vargas et al. 2017), it is important to know if and how dorsal pulvinar activity reflects spatial decisions and movement planning. Memory delay activity modulation was found in 46% of dPul units. Surprisingly, delay period responses were characterized mainly by non-space-specific suppression, rather than the spatially-specific enhancement that is more common in oculomotor thalamus and frontoparietal cortex. In dPul, suppression during the memory period was twice as frequent as enhancement, and in units showing such suppression the activity was often ramping down to a minimum around the go signal, whereas enhancement was typically sustained throughout the memory period.

The gradual decrease in firing rate during the memory period appears to be a complementary pattern to a gradual ramping up / climbing towards the saccade onset often found in frontoparietal cortex (Gnadt and Andersen 1988; Lawrence et al. 2005; Rorie et al. 2010; Premereur et al. 2011; Hwang and Andersen 2012) and in oculomotor thalamus (Watanabe and Funahashi 2004a; Tanaka and Kunimatsu 2011). Of course, the suppression during the memory delay has also been reported in these regions, but it seems to characterize only a minority of neurons (Barash et al. 1991a; Wyder et al. 2003; Watanabe and Funahashi 2004a). Hence, if we assume that the prevalent delay period activity in the frontoparietal cortex is spatially-specific excitation, the suppression in dPul implies inhibitory projections (e.g. directly or via the thalamic reticular nucleus, TRN) either from cortex to pulvinar or from pulvinar to cortex. It is important to note however that most of the suppression was not spatially-specific, unlike the cortical delay activity; hence, the prevalent patterns in dPul and cortex are not simply an inverse of each other. The non-spatial suppression might be related to a role in general alertness, task attention or non-specific movement preparation rather than specific movement planning. Unlike the “default mode” posterior cingulate neurons where the activity is suppressed starting from the onset of task-related fixation (Hayden et al. 2009), many dPul neurons decrease their firing rate below that in the initial fixation period, following a brief peripheral cue. This is interesting because the visual conditions are the same in the initial fixation and the memory delay epochs (only central fixation point is visible), but the cognitive state is different – expectation of a peripheral visual cue vs. maintenance of spatial information, motor preparation and, at the same time, attention to an upcoming foveal go signal.

Apart from mainly suppressive non-spatial signals, some dPul units (16%) also showed differential memory period responses for upcoming ipsilateral and contralateral instructed saccades, indicating potential involvement in directly linking the visual cue and the upcoming action. There was no contralateral preference of memory activity on a population level, but there was an asymmetry in the two subgroups. The contralaterally-tuned units also showed contralateral cue responses and some ramping enhancement in the end of the delay relative to the fixation baseline. The ipsilaterally-tuned units did not show matching ipsilateral cue responses or ipsilateral ramping, but a suppression for contralateral trials. For comparison, 25% of units in central thalamus showed spatially-specific, mainly contralateral delay period activity in a visually-guided delayed saccade task (which is likely to produce stronger delay period activity due to the presence of visual stimulus during the delay), and mostly congruent cue and delay period tuning (Wyder et al. 2003). An even larger fraction of MD units (53%) showed mainly contralateral and cue-congruent excitatory delay period activity (Watanabe and Funahashi 2004a). Less predominant and less contralateral delay period activity sets dPul apart from those oculomotor thalamic nuclei.

Another likely distinction of dPul from the central thalamus and MD is the degree to which these structures show prospective target selection. In rotation or bilateral target color-matching choice memory saccade tasks, many neurons in MD and the central thalamus prospectively encode the direction of the upcoming saccade, rather than visual cues (Watanabe and Funahashi 2004b; Wyder et al. 2004). In contrast, memory period and pre-saccadic activity in spatially selective dPul neurons did not signal the upcoming movement direction during bilateral cue free choice trials. Choice selectivity analysis indicated that target selection was not reflected in the epochs before the saccade onset, because only a very small amount of units significantly preferred the same hemifield in both instructed and choice trials. While it is worth noting that the free choice where both targets are “correct” differs from the above tasks because they unequivocally specify the correct target, it stands to reason that oculomotor thalamus would exhibit a prospective target selection also in the free choice task, as was demonstrated in the frontoparietal cortex (Coe et al. 2002; Watanabe and Funahashi 2007). The lack of congruent (to instructed trials) choice selectivity in dPul suggests that the observed spatial modulation in the instructed trials reflects the retrospective visual processing rather than the spatial decision for the upcoming movement (Lindner et al. 2010; Lundqvist et al. 2018).

The finding that the dPul delay period activity does not encode the upcoming choice might appear to challenge the underlying notion that the pulvinar activity during the memory delay period is directly involved in decision making, derived from the inactivation studies (Wilke, Turchi, et al. 2010; Wilke et al. 2013). On the other hand, the lack of choice encoding is consistent with our recent findings showing the lack of the choice bias induced by the transient dPul microstimulation during the memory delay, as compared to strong choice effect in the pre-saccadic epoch of the visually-guided task (Dominguez-Vargas et al. 2017). But we emphasize that neither of those findings invalidate the results showing a contralesional choice deficit during memory saccades due to dPul inactivation. Such choice biases are likely caused by disrupting contralesional visual and less prominent pre/peri-saccadic activity, as well as, more hypothetically, the across-trial changes in post-saccadic confidence or feedback evaluation affecting future target selection. Furthermore, since such dPul activity is transmitted to the cortex, the dPul inactivation is expected to strongly affect cortical contralesional spatial representations and ensuing visuomotor decisions (Wilke, Kagan, et al. 2010; Jaramillo et al. 2019; Klink et al. 2021).

Furthermore, and not mutually exclusive to encoding visuomotor decision and motor planning, it has been shown that both dorsal and ventral lateral pulvinar causally contribute to orienting spatial attention (Petersen et al. 1987; Desimone et al. 1990; Zhou, Schafer, et al. 2016). In terms of the underlying neuronal signals, two studies in the ventral lateral pulvinar (Saalmann et al. 2012; Zhou, Schafer, et al. 2016) and one recent study in the medial dorsal pulvinar (Fiebelkorn et al. 2019) reported slight but significantly enhanced persistent activity during sustained attention to a visible stimulus in the receptive field. It is hence conceivable that a presence of a visual stimulus, which is attended to or is a goal of an upcoming movement, is required for the pulvinar to exhibit the delay period enhancement and spatial selectivity.

Overall, a considerable response modulation during the delay period notwithstanding, our results strengthen a prior conclusion (Bender and Baizer 1990) that the primary oculomotor role of the pulvinar is not in neural events leading to prospective saccade preparation and generation, at least in the absence of a visible goal (i.e. as in the memory saccade task). Rather, dPul is mainly involved in integrating consequences of saccadic eye movements (saccade direction, amplitude, time of occurrence) with visual processing.

### Future directions

The importance of systematic characterization of basic visuomotor properties of a complex and still not well understood structure such as the dorsal pulvinar cannot not be underestimated. At the same time, the insights gained with simple oculomotor tasks like visually- and memory-guided saccades are inherently limited, given the diversity of pulvinar responses and its proposed involvement in multiple cognitive processes. More elaborate tasks are needed to further evaluate and dissociate involvement of dorsal pulvinar neurons in visual processing, spatial target selection, attentional and motivational factors. One way is to utilize paradigms that dissociate the visual input and the motor output, e.g. anti-saccade or rotated saccade tasks (Munoz and Everling 2004; Watanabe and Funahashi 2004b; Kunimatsu and Tanaka 2010). Second, the tasks that combine immediate stimulus information with a decision urgency, such as the compelled saccade task (Costello et al. 2015) or a difficult perceptual discrimination under a time pressure, might evoke more prospective encoding in the pulvinar neurons. Third, choice tasks with an explicit “correct” response e.g. color matching target/distractor choice task (Wyder et al. 2004), or reward modulation where different color targets provide a different amount of reward (Wilke et al. 2013), may inform about the encoding of behavioral salience and selected action.

Beyond the question of target selection in the memory-guided oculomotor task, dPul might be more involved in guiding other movements such as hand actions, as suggested by the effect of pulvinar inactivation or lesions on target selection via reaching and grasping, and hand-specific deficits (Wilke, Turchi, et al. 2010; Wilke et al. 2018), as well as its strong connectivity to the medial posterior parietal areas (Cappe et al. 2007; Gamberini et al. 2021, 2021). Furthermore, given that many dPul units show gain-field like properties or other types of postural modulations by the position of the eyes in the orbit (Schneider et al. 2019), it remains important to assess if pulvinar is causally involved in spatial reference frame transformations and eye-hand coordination. Rodent studies also suggests that the medial part of the lateral posterior nucleus, the homolog of the primate pulvinar, is connected to amygdala, striatum and frontoparietal cortex and integrates sensorimotor multimodal information, providing an evolutionary perspective on the role of the pulvinar in coordination of perception and body movements (Zhou et al. 2017; Leow et al. 2022).

## Conclusions

The response timing and tuning suggest that the dorsal pulvinar contributes to spatial actions largely by supporting visuomotor processing around visual cues and saccades. Many neurons show a conjunction of at least two of the three components - visual, delay and/or motor responses. But properties of most individual neurons are inconsistent with prospective transformation of spatial goals into actions during the working memory saccade task. These findings do not rule out more direct involvement of the dorsal pulvinar at the level of single neurons to prospective planning and decision in other contexts. Furthermore, beyond the single neuron perspective, the dorsal pulvinar is well positioned to mediate the visuomotor transformations and spatial decisions at the level of neuronal ensembles comprising groups of neurons encoding different task components, locally and via the connectivity to the cortex.

## Authors contributions

LS, AUDV, MW and IK designed the experiments, AUDV and LG collected the data and performed initial analyses, LS analyzed the data and created the figures, IK edited the figures, LS, MW and IK wrote the paper.

## Acknowledgements

We thank Ira Panolias, Daniela Lazzarini, Sina Plümer, Klaus Heisig, Leonore Burchardt, and Dirk Prüße for technical support. We thank Stefan Treue, Alexander Gail, Hansjörg Scherberger, members of the Decision and Awareness Group, Sensorimotor Group and the Cognitive Neuroscience Laboratory for helpful discussions.

