## Supplementary Material for "Visual, delay and oculomotor timing and tuning in macaque dorsal pulvinar during instructed and free choice memory saccades"

6 Supplementary Figures

1 Supplementary Table

List of Abbreviations

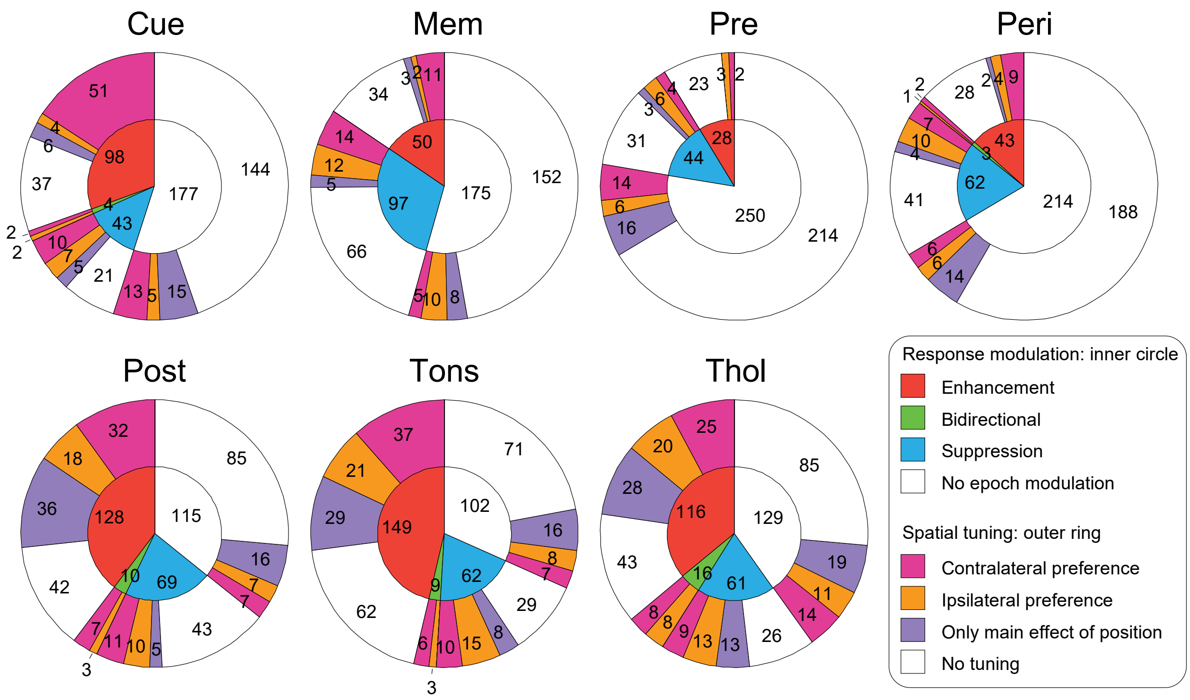


**Supplementary Figure S1. Relationship of spatial tuning and response modulation per epoch.** Related to **Figure 3**. Spatial tuning and response modulation of all 322 units recorded during the memory-guided saccade task. Spatial preference and non-space specific modulation in seven epochs: fixation hold (Fhol), cue onset (Cue), memory (Mem), pre-saccadic (Pre), Peri-saccadic (Peri), Post-saccadic (Post), target onset (Tons), and target hold (Thol), see **Materials and Methods**). In each plot, sectors of the inner circle display the number of units that showed statistically-significant enhancement or suppression in the respective epoch as compared to the respective baseline epoch: units showing only enhancement (red), only suppression (blue), enhancement for one hemifield and suppression for the other (green), or no enhancement nor suppression (white). The outer sectors display the number of units that preferred the contralateral hemifield (purple), the ipsilateral hemifield (orange), did not prefer either hemifield but showed a main effect of position in a one-way ANOVA (purple), or were not tuned (white), for each of the four response modulation subsets.

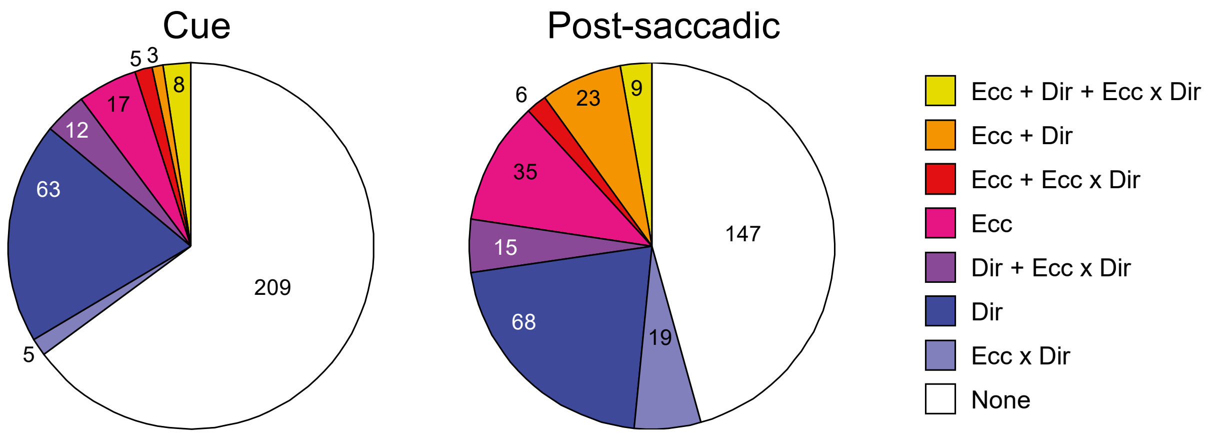


**Supplementary Figure S2. Direction (Dir) and absolute eccentricity (Ecc) dependence.** Related to **Figure 4**. Results of two-way ANOVA on firing rates, with main factors direction (one of six target directions, *cf.* **Figure 1A** inset) and absolute eccentricity (12° or 24°).

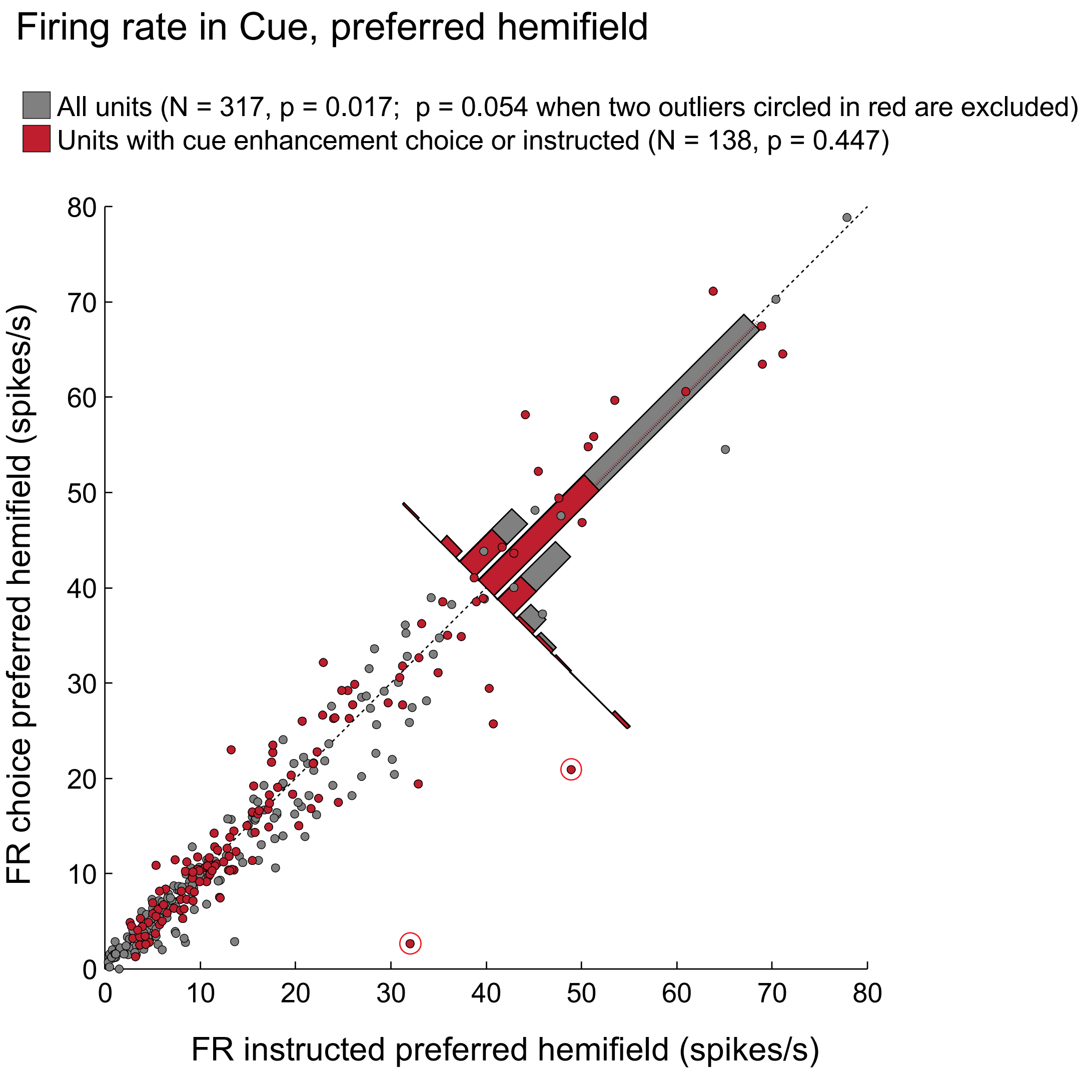


**Supplementary Figure S3. Instructed (unilateral single target) vs. choice (bilateral double targets) cue response.** Related to **Figure 4**. Scatter plot for the cue response firing rate in the choice condition when the preferred hemifield was selected vs. the preferred instructed condition. Gray or red dots – all units with 4 or more trials for each hemifield in both instructed/unilateral and choice/bilateral conditions, N=317; red dots – units with significant cue enhancement in either instructed or choice condition, N=138. Across all units, the instructed response was stronger than the choice (paired t-test, p = 0.017, but without the two circled "outlier" units only approaching significance at p = 0.054); for units with significant cue enhancement, there was no difference (p=0.477). Across all units with a significant enhancement RF zone, units showing spatial summation had larger RFs (bilateral>unilateral: N=54, 16.9±6.5°) than units showing suppression (unilateral>bilateral: N=82, 14.6±6.4°; p=0.046, unpaired t-test).

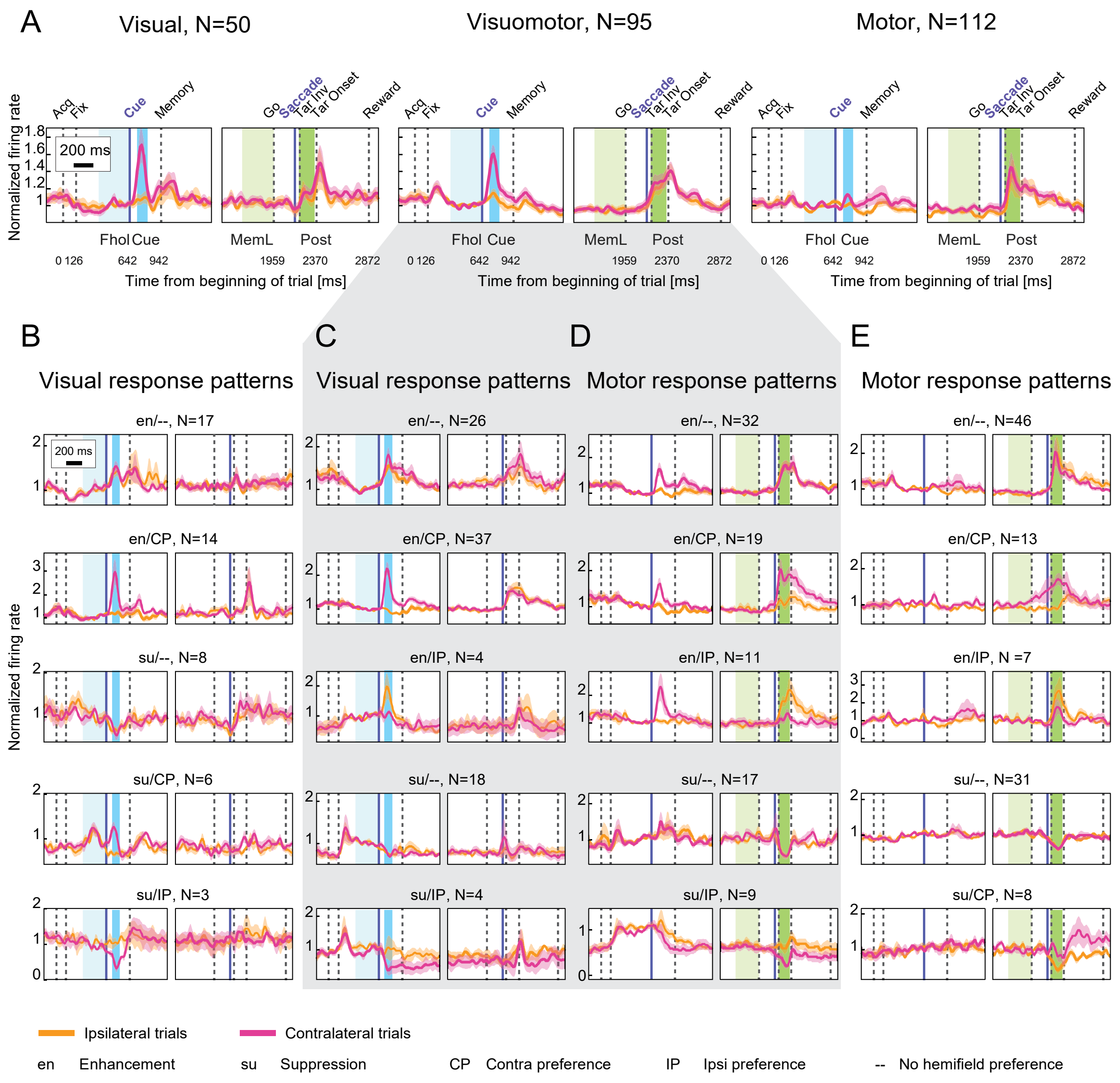


**Supplementary Figure S4. Population responses of visual, visuomotor, and motor units.** Related to **Table 1**. Vertical lines mark events, and colored areas relevant epochs for categorization. **A:** Population response of the three categories for contralateral (magenta) and ipsilateral (orange) trials: visual (enhancement or suppression in Cue relative to Fhol, but not in Post), motor (enhancement or suppression in Post relative to Mem, but not in Cue) and visuomotor units (enhancement or suppression in both epochs). **B:** Subpopulations of visual cells grouped by most common visual response patterns. **C:** Subpopulations of visuomotor cells grouped by most common visual response patterns. **D:** Subpopulations of visuomotor cells grouped by most common motor response patterns. **E:** Subpopulations of motor cells grouped by most common motor response patterns; "en/--": enhancement without hemifield preference; "su/--": suppression without hemifield preference; "en/CP": enhancement and contralateral preference; "en/IP": enhancement and ipsilateral preference; "su/CP": suppression and contralateral preference; "su/IP": suppression and ipsilateral preference.

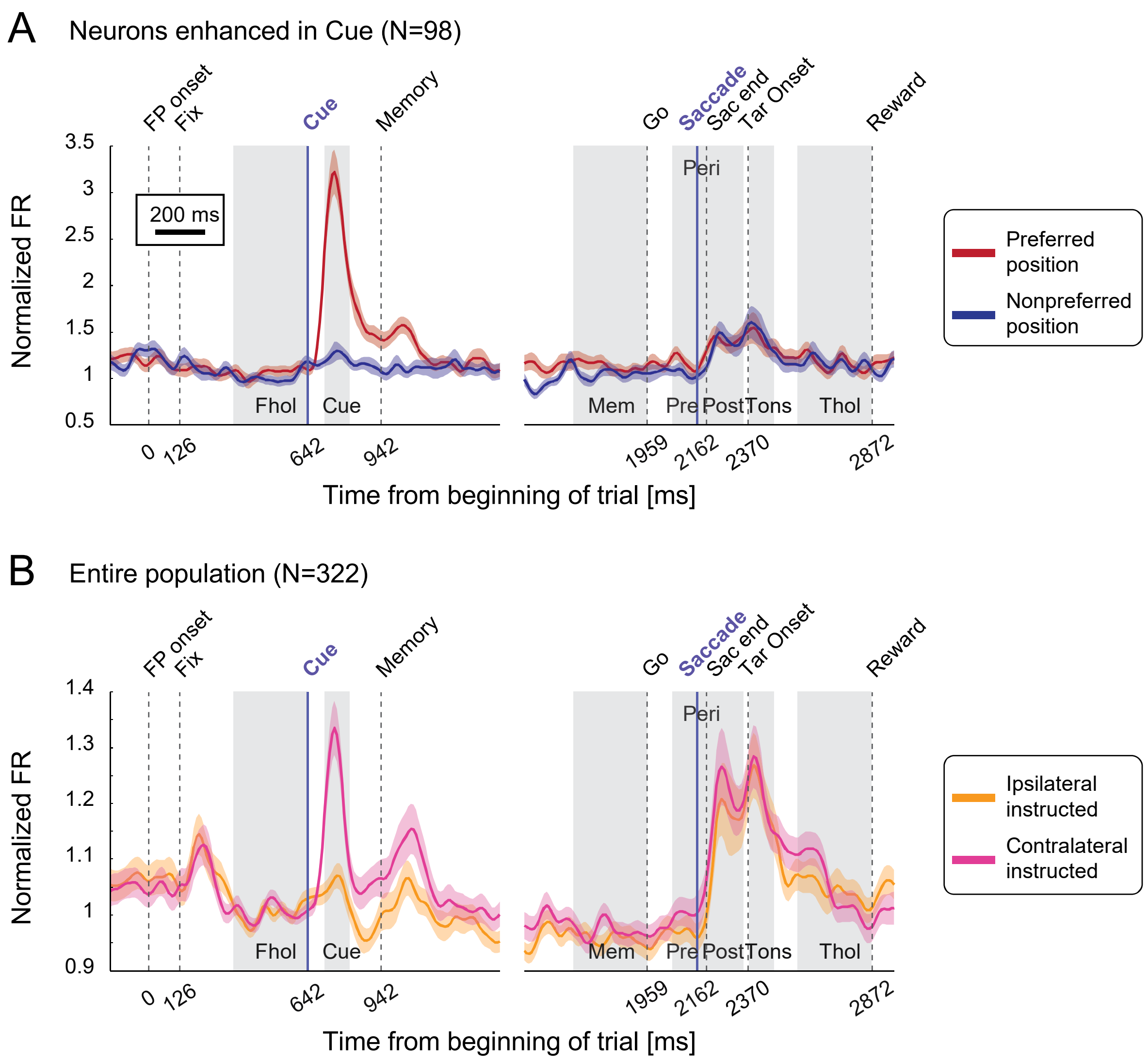


**Supplementary Figure S5. Population activity.** Related to **Figure 8G-H.** Population responses, normalized by dividing by the activity in the fixation hold epoch. **A:** Population average for units showing enhancement in the cue epoch relative to fixation hold epoch (Fhol), for the preferred target position for the cue epoch (dark red) and the opposite target position (nonpreferred - dark blue). **B:** Population average across the entire population for contralateral (magenta) and ipsilateral (orange) trials.

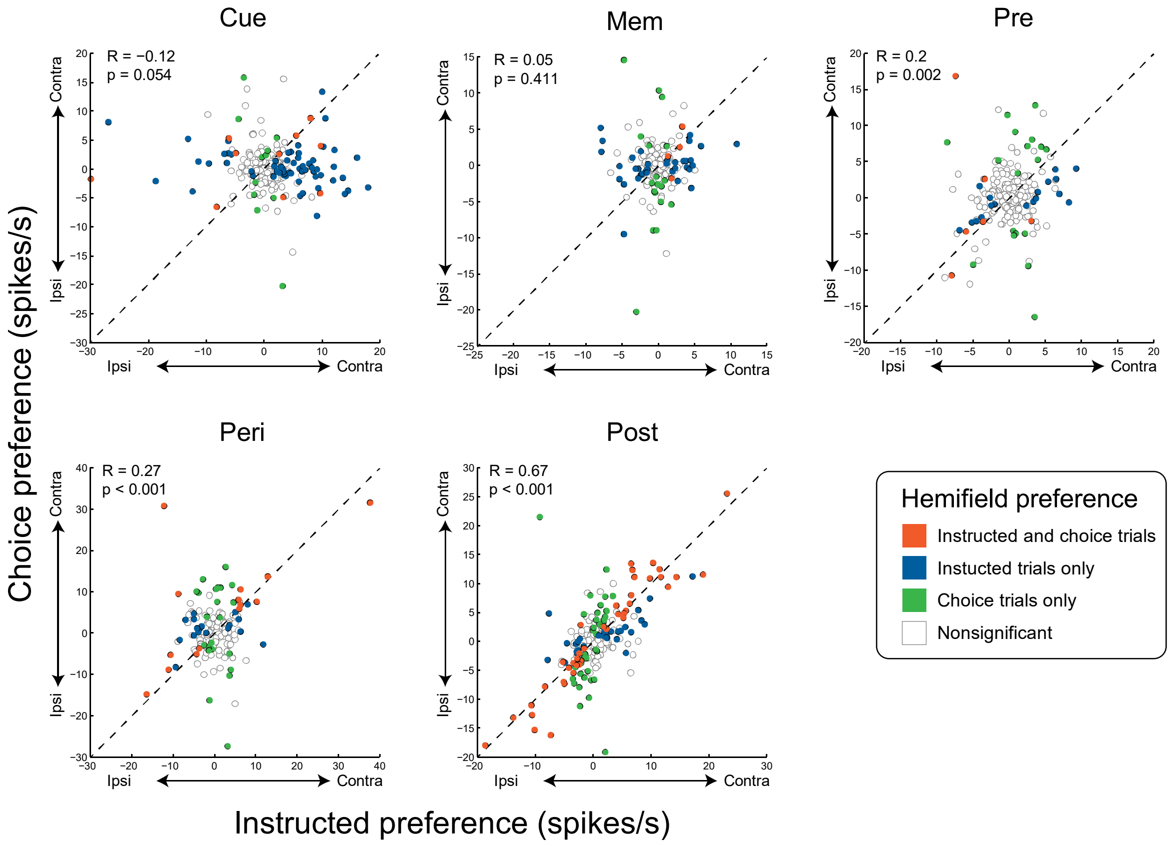


**Supplementary Figure S6. Analysis of choice preference in single units**. Related to **Figure 9C**. Scatter plots comparing firing rate hemifield preference ($FR_{contra}$ – $FR_{ipsi}$) in instructed and choice trials, for six epochs: cue onset (Cue), memory (Mem), pre-saccadic (Pre), peri-saccadic (Pre) and post-saccadic (Post). R and p values for Pearson's correlations between hemifield preference in instructed and choice trials are shown. Four units were excluded from this analysis because less than 4 choice trials for one of the two hemifields were recorded, resulting in remaining N=245 units. Note that since the classification to multi-units and single units was performed separately per each recording block (120 trials), in few cases (N=14) the same unit was considered a single unit in some block(s) but a multi-unit in other block(s) if the isolation became less or more clear. In such cases, data from all blocks were included in **Figure 9C** but only data from blocks with better separation in this figure – hence the individual data points in this figure are not always a subset of points in **Figure 9C**.

**Supplementary Table S1. Saccade reaction times (RT), durations and maximum velocities.** All values are means across sessions ± standard error, separately per task, monkey and instructed or choice condition, either per hemifield (Left/Right) or both hemifields combined (All).

| **Task** | **Monkey** | | **RT [ms]** | | | **Duration [ms]** | | | **Velocity [°/s]** | | |
|---|---|---|---|---|---|---|---|---|---|---|---|
| | | | All | Left | Right | All | Left | Right | All | Left | Right |
| **Memory guided** | C | Instructed | 211±1 | 207±2 | 217±1 | 46.8 | 47.5±0.2 | 47.6±0.3 | 594±3 | 605±5 | 581±6 |
| | | Choice | 206±1 | 204±1 | 209±2 | 46.4 | 44.5±0.5 | 50.7±0.8 | 602±3 | 570±4 | 645±9 |
| | L | Instructed | 191±1 | 193±2 | 189±1 | 44.9 | 44.3±0.2 | 46.0±0.3 | 640±4 | 671±5 | 598±4 |
| | | Choice | 191±1 | 195±2 | 186±1 | 45.4 | 47.8±0.6 | 43.0±0.6 | 648±6 | 726±7 | 558±6 |
| **Visually guided** | C | Instructed | 177±1 | 180±1 | 174±1 | 43.2 | 43.4±0.2 | 43.6±0.3 | 681±3 | 706±5 | 651±7 |
| | | Choice | 172±1 | 174±2 | 172±1 | 42.8 | 41.9±0.3 | 43.7±0.3 | 675±4 | 713±8 | 653±7 |
| | L | Instructed | 163±1 | 166±1 | 160±1 | 41.8 | 41.7±0.1 | 42.0±0.2 | 707±3 | 744±3 | 663±4 |
| | | Choice | 169±2 | 177±2 | 166±1 | 41.9 | 42.1±0.3 | 41.6±0.3 | 693±5 | 758±22 | 650±4 |

### List of Abbreviations

| | |
|---|---|
| ANOVA | Analysis of variance |
| CI | Contralaterality index |
| FR | Firing rate |
| LED | Light-emitting diode |
| MATLAB | Matrix laboratory |
| MRI | Magnetic resonance imaging |
| PEEK | Polyetheretherketone |
| RARE | Rapid acquisition with relaxation enhancement |
| PSTH | Peri-stimulus time histogram |
| RF | Receptive / response field |
| SD | Standard deviation |
| SE | Standard error |

**Anatomical labels**

| | |
|---|---|
| dlPFC | Dorsolateral prefrontal cortex |
| dPul | Dorsal pulvinar |
| FEF | Frontal eye field |
| IPul | Inferior pulvinar |
| LIP | Lateral intraparietal cortex |
| LPul | Lateral pulvinar |
| MD | Mediodorsal nucelus |
| MPul | Medial pulvinar |
| PLdm | Dorsomedial lateral pulvinar |
| PLvl | Ventrolateral lateral pulvinar |
| PPC | Posterior parietal cortex |
| SC | Superior colliculus |
| TPO | Temporo-parieto-occipital area |
| VIP | Ventral intraparietal cortex |
| vPul | Ventral pulvinar |
